# Minute-scale control of ubiquitin-mediated degradation reveals dynamics of bacterial secreted effector-functions

**DOI:** 10.1101/2025.11.19.688170

**Authors:** Haolin Zhang, Yongxia Guo, Bikash Adhikari, Nevenka Dudvarski-Stankovic, Elmar Wolf, Thomas Rudel

## Abstract

Precise temporal control of protein abundance is essential for dissecting dynamic cellular processes. While degron-based systems enable rapid protein depletion in eukaryotic cells, comparable tools are lacking for bacterial effectors delivered into host cells during infection. Here, we establish AIDE (Auxin-Inducible Degradation of Effectors), a host-directed degradation platform that harnesses the ubiquitin–proteasome system to selectively eliminate secreted bacterial proteins, including membrane-integrated effectors. By integrating a minimal auxin-inducible degron (AID) tag into effector genes, AIDE enables rapid, reversible, and spatially confined degradation while preserving native expression and secretion. Applied to *Chlamydia trachomatis*, AIDE revealed that the membrane-integrated deubiquitinase Cdu1 suppresses autophagy early and later promotes developmental transitions, whereas the integral membrane fusogen IncA is continuously required for inclusion integrity. This AIDE platform provides minute-scale, spatiotemporal control over bacterial effector activity, offering a broadly applicable framework for dissecting virulence mechanisms and host–pathogen interactions across diverse secretion-dependent pathogens.

## Introduction

Bacterial infections are the second leading cause of death worldwide ^1^. Many pathogenic bacteria cause diseases by delivering proteins into host cells via specialized secretion systems. Among these, the bacterial type III secretion system (T3SS) is a protein transport apparatus used by numerous Gram-negative bacteria to inject effector proteins directly into host cells. Also known as an injectisome, the T3SS functions like a “nanosyringe” enabling bacteria to deliver proteins straight into the host cytoplasm or organelles, bypassing the extracellular space ^2, 3^. These effector proteins manipulate host cell processes to facilitate infection ^4, 5^. A functional injectisome is critical for the pathogenicity of many important bacterial pathogens, including agents of foodborne and waterborne disease such as *enteropathogenic Escherichia coli*, *Salmonella enterica* serovar Typhimurium, and *Shigella dysenteriae*; insect-borne and zoonotic pathogens such as *Rickettsia spp.* or *Yersinia pestis*; nosocomial pathogens like *Pseudomonas aeruginosa*; and sexually transmitted pathogens such as *Chlamydia trachomatis* ^2, 6^. T3SSs are highly dynamic structures that can inject their substrates into host cells within minutes ^7^. Many of the secreted effector proteins have enzymatic or structural functions, which have traditionally been studied either by loss-of-function approaches in the pathogenic bacteria or by overexpression in the host cell of interest ^8^. Both strategies have limitations: loss-of-function screens are slow and carry the risk of compensatory events arising during mutant selection, while overexpression in host cells may produce artifacts. Similarly, overexpression of bacterial effectors in host cells often leads to their mislocalization to inappropriate cellular compartments. Moreover, unphysiological levels of effectors can create artifacts by disrupting the balance of host cell processes. For many obligate intracellular bacteria, effectors may be essential for bacterial survival or replication, meaning that gene knockouts can compromise the viability of the pathogen ^9, 10^.

A strategy that could overcome these limitations is the rapid and controllable degradation of secreted bacterial effectors within host cells. In eukaryotes, targeted protein degradation (TPD) has been developed as a therapeutic approach to eliminate specific proteins ^11, 12^. A prominent example is the use of bifunctional small molecules, known as PROTACs, which recruit target proteins to E3 ubiquitin ligases for proteasomal degradation. However, PROTACs development relies on the presence of druggable domains within the target protein, requires extensive medicinal chemistry optimization, and often fails ^13, 14^. An alternative approach is genetic degron systems, which fuse ligand-responsive tags to proteins of interest, enabling controlled degradation ^15–18^.

The auxin-inducible degron (AID) system exploits the plant F-box receptor TIR1, a substrate adaptor of the SCF E3 ligase complex (Skp1-Cullin1-Rbx1-TIR1). Upon auxin addition, the hormone binds to the leucine-rich repeat pocket of TIR1, promoting formation of a TIR1-auxin-AID ternary complex with the degron-tagged substrate. This interaction triggers K48-linked polyubiquitination and subsequent 26S proteasome-mediated degradation of the target protein. In most cases, tagged proteins are efficiently depleted within approximately 20-40 minutes of auxin treatment ^16^. The system was recently upgraded to the second-generation AID system (AID2), which uses engineered OsTIR1(F74G) and the synthetic auxin analog 5-Ph-IAA in a “bump-and-hole” strategy. This modification virtually eliminates background degradation and dramatically increases ligand sensitivity, while further accelerating degradation kinetics ^19^. With high specificity and efficiency, AID2 allows precise, reversible protein depletion, making it ideal for studying dynamic cellular processes.

We developed a platform for Auxin-Inducible Degradation of Effectors (AIDE). As a proof of principle, we focused on *Chlamydia trachomatis*, an obligate intracellular pathogen and the most common cause of sexually transmitted infections in humans ^20^. *C. trachomatis* replicates within a membrane-bound compartment in the host cell, known as an inclusion, following a complex developmental cycle. Infectious elementary bodies (EBs) enter host cells and differentiate into replicative reticulate bodies (RBs). After several rounds of replication, RBs convert back into EBs, which exit the host cell to initiate new rounds of infection ^21, 22^. T3SS effectors are essential at all stages of chlamydial development, contributing to host cell entry ^21, 23^, inclusion formation and stabilization ^24–26^, and the manipulation of multiple host signaling pathways to acquire nutrients and evade cell-autonomous defense mechanisms ^21, 27^. The chlamydial T3SS effector repertoire is estimated to comprise over 100 proteins, yet only a few have been functionally characterized ^28, 29^. Among these effectors, inclusion membrane proteins (Incs) are critical for pathogen survival. These proteins are embedded into the inclusion membrane and support the inclusion integrity and the development of *Chlamydia* by directly manipulating numerous host cell pathways.

The inducible direct degradation of effectors as we planned to establish in the AIDE system allows investigations into the dynamics and function of secreted proteins that are not approachable by silencing technologies. These technologies work by inducible expression of regulatory RNAs ^30^ or the inducible expression of guide RNAs in CRISPR-expressing *Chlamydia* ^31, 32^ to interfere with protein translation. Since these systems work by preventing new synthesis of proteins, they are used to determine the effect of inhibiting translation in general since a defined depletion of a protein in a short period of time to access the dynamics of function is not possible ^33^. One reason for these limitations is the extremely different half-life of proteins, particularly the half-life of membrane proteins ranges from several hours up to days ^34^.

One of the Incs from *Chlamydia* particularly interesting to access by AIDE is the Chlamydia Deubiquitinase 1 (Cdu1) which interferes with inclusion ubiquitination, autophagy signaling and thereby supports recruitment of Golgi-derived vesicles to the inclusion to support nutrient acquisition ^35–37^. Cdu1 has two distinct enzymatic activities. Its deubiquitinase function removes ubiquitin tags, while its acetyltransferase activity acetylates itself and chlamydial effectors (e.g., IpaM, InaC) at lysin residues to protect them from ubiquitination^37^. Although both the deubiquitinating and acetyltransferase activities of Cdu1 have been demonstrated with defined mutants, the dynamics of these functions during infection, such as whether the inclusion remains ubiquitin-free for a defined period in the absence of Cdu1, have yet to be elucidated.

Another Inc protein whose temporal dynamics remains to be elucidated is IncA. By mimicking host SNARE proteins, IncA mediates inclusion fusion and prevents lysosomal degradation by competitively inhibiting interactions with host proteins such as Vamp3 and Vamp8 ^38, 39^. A key unresolved question central to the understanding IncA’s role is whether IncA acts exclusively to initiate inclusion fusion or is also required to maintain the stability of fused inclusions over time.

Here, we implemented the AIDE system to harness the host degradation machinery for the selective clearance of bacterial effectors at the host-Chlamydia interface, generating Ctr-AIDE. Using this platform, we dissected the temporal functions of Cdu1 and IncA, revealing their phase-specific roles in evasion of autophagy signaling, developmental transitions, and inclusion stability. This study therefore provides a blueprint for the use of AIDE as a tool to analyze the dynamic functions of secreted effectors during bacterial infection.

## Results

### A Genome-Integrated Platform for Conditional Protein Depletion

To establish the AIDE system in *Chlamydia trachomatis* (*Ctr*), we designed a strategy to integrate degron sequences into chlamydial target effector proteins (Ctr-AIDE). Central to this system is a 7-kDa mAID degron, fused to target effectors and recognized by the mutant host F-box receptor OsTIR1(F74G), which forms the SCF E3 ligase complex and thus directs K48-linked ubiquitination upon addition of the synthetic auxin 5-Ph-IAA. To enable precise degron integration into *Ctr*, we combined AID2 with the FRAEM (Fluorescence-Reported Allelic Exchange Mutagenesis) system, a homologous recombination-based method for seamless genome editing ^40^. This integration yielded Ctr-AIDE, a platform for rapid and reversible depletion of secreted effectors (Fig. 1A).

**Figure 1.**
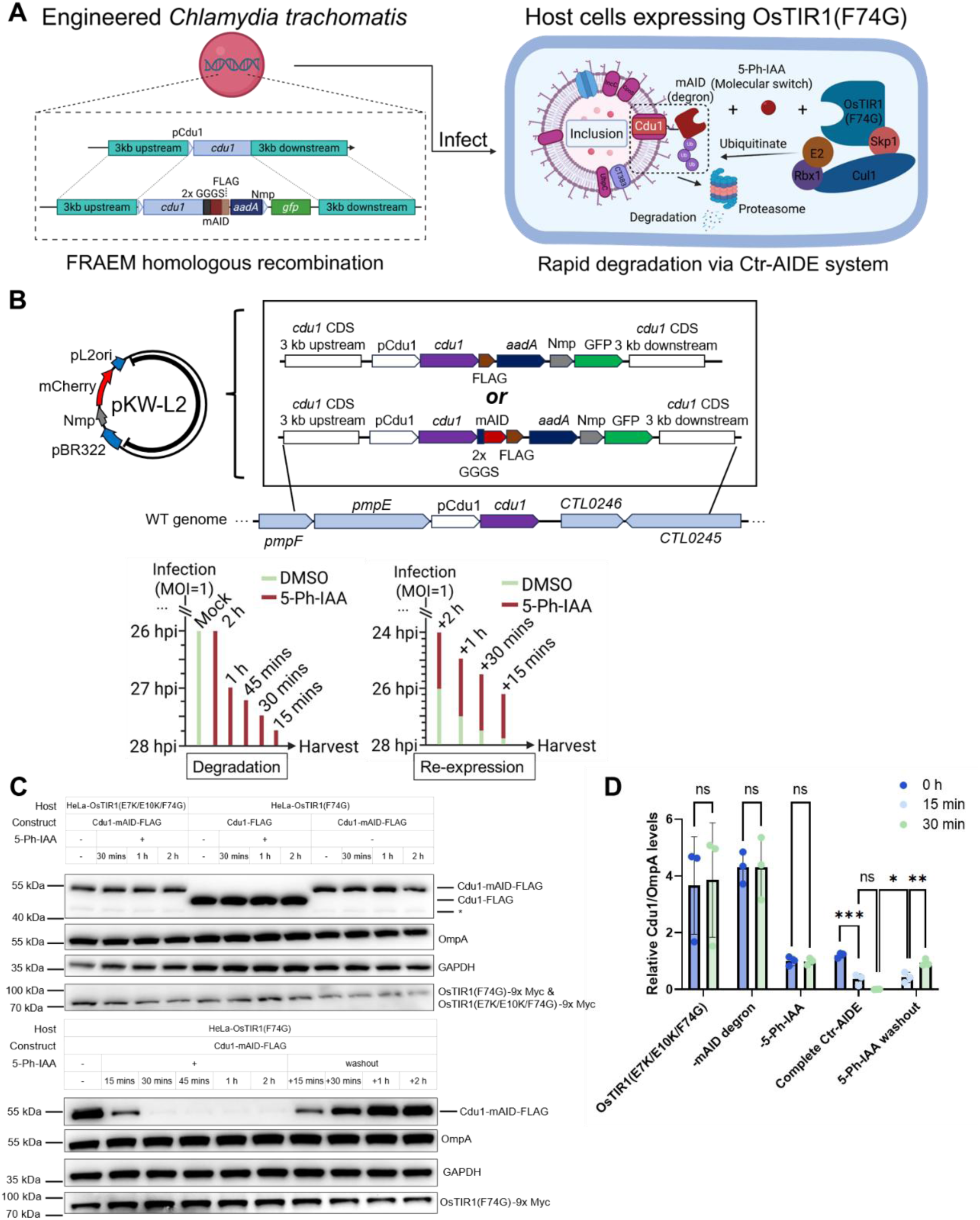
AID2 system enables temporal control of Cdu1 expression in *Chlamydia*-infected HeLa cells. (A) Schematic of Ctr-AIDE strategy, combining the FRAEM genome-editing platform (left) with the AID2 TPD system (right). Cdu1 is shown as proof of concept. (B) *Top*: Plasmid constructs for FRAEM-mediated homologous recombination in *Ctr* serovar L2. The targeted genomic region is marked by lines. pL2ori represents the putative chlamydial plasmid ori, and pBR322 (Ori) is included to facilitate cloning in *E. coli*. pCdu1, native Cdu1 promoter; Nmp, *Neisseria meningitidis* promoter; *aadA*, spectinomycin resistance gene; *Bottom*: Ctr-AIDE degradation/re-expression strategy. Vertical bars indicate treatment with DMSO (control) or 5-Ph-IAA (1 µM). All samples were harvested at 28 hours post-infection (hpi) for downstream analysis. (C) Ctr-AID-mediated regulation of Cdu1-mAID. Cells were infected with indicated strains at MOI=1. Cdu1 (anti-FLAG) and OsTIR1 mutants (anti-Myc) levels were monitored, with Chlamydial Major Outer Membrane Protein (OmpA) as a loading control and GAPDH as a host cell control. (D) Quantification of Cdu1 degradation. FLAG signal (normalized to OmpA, mean ± SD) from three independent immunoblot (one shown in C). Statistical significance determined by two-tailed paired t-test (****p < 0.0001; n.s., not significant). Controls: OsTIR1(E7K/E10K/F74G), -mAID degron, -5-Ph-IAA: controls with dysfunctional E3 ligase OsTIR1(F74G), lacking mAID tagging, and lacking 5-Ph-IAA treatment, respectively. All experiments were performed ≥3 times with consistent results.

For several reasons, we selected the chlamydial deubiquitinase Cdu1 to establish the Ctr-AIDE strategy: (i) Cdu1 is a highly active K48 deubiquitinase that upon AIDE (that ubiquitinated K48 to induce degradation) could lower the efficiency of degradation; (ii) The auto-acetylation of Cdu1 at lysin residues has been implicated in ubiquitination resistance of the protein ^37^; (iii) As a membrane integrated protein, degradation of Cdu1 is expected to be particularly challenging; (iv) Rapid Cdu1 protein degradation and re-expression could be a way to shed light on the effect of the deubiquitinating and acetyltransferase activities of Cdu1 in ongoing infections. To generate the Cdu1 degradation construct, a knock-in cassette was designed in the suicide plasmid pKW-L2 ^40^, consisting of (i) 3 kb homology sequences flanking the *cdu1* locus; (ii) an mAID-FLAG sequence fused to the C-terminus of Cdu1 via a flexible linker (2x GGGS), ensuring cytosolic accessibility given that the Cdu1 C-terminus faces the host cytosol ^35^; and (iii) a GFP selection cassette with spectinomycin resistance gene (aadA-GFP, Fig. 1B). This design enables allelic replacement of endogenous *cdu1* with the mAID-FLAG-tagged variant, ensuring native expression regulation and eliminating reliance on plasmid-based expression. *Ctr* recombinants were initially screened for loss of the plasmid pKW-L2 (RFP^-^) and the expression of GFP indicative of successful recombination. In addition, mAID tagging was confirmed by anti-FLAG immunoblotting (Fig. S1A). A matched Cdu1-FLAG control strain that lacks the mAID degron was constructed in parallel for comparative analysis (Fig. S1A).

We established stable HeLa cell lines expressing either functional OsTIR1(F74G) or catalytically impaired OsTIR1(E7K/E10K/F74G) ^41^, and validated AID2 functionality by mAID-GFP degradation. The impaired OsTIR1(E7K/E10K/F74G) mutant served as a reliable degradation-negative control (Fig. S2A-C). Using this system, we investigated Ctr-AIDE-mediated depletion of the effector Cdu1. No K48 ubiquitin was detected on the inclusions in the Ctr-AIDE strain indicating that Cdu1-mAID functions in protecting the inclusion from ubiquitination. Strong K48 ubiquitination was visible on the inclusion already 5 minutes after addition of the 5-Ph-IAA inducer (Fig. S3) suggesting that OsTIR1(F74G) is recruited to Cdu1-mAID and induces its ubiquitination. Surprisingly, strong degradation of Cdu1-mAID was visible already 15 minutes after addition of the 5-Ph-IAA inducer and degradation was complete within 30 minutes. Degradation depended on the mAID tag since Cdu1 remained stable in (i) cells expressing the catalytically impaired OsTIR1(E7K/E10K/F74G) variant ^41^, (ii) strains expressing non-degron Cdu1-FLAG constructs, and (iii) untreated conditions. Importantly, degradation was fully reversible, with Cdu1-mAID re-accumulating partially within 15 minutes and completely within 2 hours following 5-Ph-IAA washout after 2 hours of treatment (Fig. 1B-1D). Immunofluorescence microscopy confirmed rapid loss of inclusion membrane-localized Cdu1 after 5-Ph-IAA treatment and its restoration upon 5-Ph-IAA washout (Fig. 2A). Spatial resolution was further validated by 4 x expansion microscopy ^42^, which confirmed specific depletion of inclusion membrane-associated Cdu1 while pools in the chlamydial cytoplasm remained unaffected, highlighting the system’s spatial precision (Fig. 2B). To assess target specificity of Ctr-AIDE for mAID-tagged effector, we monitored the untagged inclusion membrane protein IncA during rapid Cdu1 depletion. Immunoblotting showed that IncA expression remained unchanged while Cdu1-mAID was rapidly degraded (Fig. S4), indicating that Ctr-AIDE selectively degrades mAID-tagged proteins without affecting untagged effectors.

**Figure 2.**
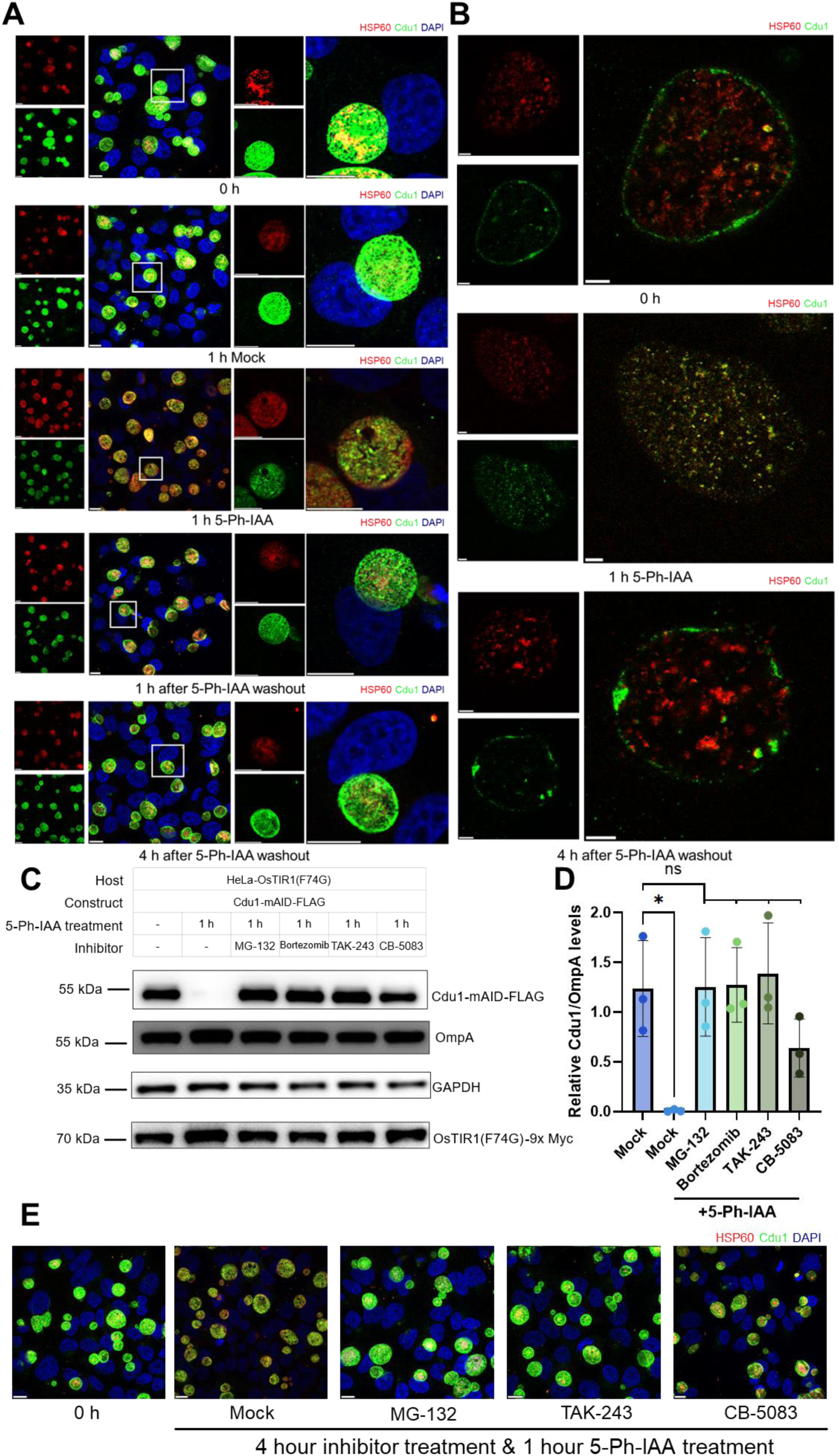
AID2 system enables spatial regulation of Cdu1 expression in *Chlamydia*-infected HeLa cells and depends on an intact ubiquitin-proteasome pathway. (A) Immunofluorescence microscopy confirms Cdu1 depletion dynamics. Cdu1 (green, anti-FLAG), chlamydial inclusion (Hsp60,red), and DNA (DAPI, blue). Scale bar = 10 µm. (B) Expansion microscopy (4× expansion) resolving Cdu1 localization patterns under undegraded and degraded conditions. Scale bar = 10 µm. (C) Cdu1 degradation depends on ubiquitin-proteasome and p97 pathways. Host cells were pretreated with inhibitors for 3 hours (including 1 hour during 5-Ph-IAA treatment) with pan-E1 (TAK-243, 1 µM), proteasome (MG-132, 10 µM; bortezomib, 1 µM), or p97 (CB-5083, 10 µM) inhibitors. Degradation was blocked despite 1-hour 5-Ph-IAA exposure. (D) Quantification of inhibitor effects on Cdu1 degradation. FLAG levels (normalized to OmpA, mean ± SD) from three independent immunoblot replicates (one representative in C). Significance assessed by one-way ANOVA with Tukey Multiple comparisons test (*p < 0.05; n.s., not significant). (E) Immunofluorescence validation of inhibitors. Inclusion-localized Cdu1 signal persists in treated cultures. Scale bar = 10 µm. All experiments were replicated ≥3 times with consistent results.

Mechanistic dissection confirmed that Cdu1-mAID degradation depends on the host ubiquitin-proteasome pathways: degradation was blocked by the inhibition of the central Ubiquitin-activating enzyme 1 (TAK-243), and the proteasomal activity (MG-132, bortezomib). Interestingly, inhibition of the p97 ATPase with CB-5083 blocks Cdu1-mAID degradation. p97 functions by extracting misfolded or damaged proteins from membranes, a process triggered by ubiquitination ^43^, suggesting that degradation of inclusion membrane proteins involves p97, potentially through extraction of membrane proteins from the inclusion prior to proteasomal degradation. (Fig. 2C-2E).

To assess the functionality of Ctr-AIDE in different cell lines, Cdu1 degradation was tested in A-375 (melanoma), HCT 116 (colon carcinoma), and U-2 OS (osteosarcoma). All lines exhibited degradation kinetics comparable to HeLa (cervical carcinoma) cells, with HeLa showing the highest proteasomal degradation efficiency (Fig. S5). These results confirm that Ctr-AIDE functions robustly across diverse host cell types, with degradation fidelity dependent on the host ubiquitin-proteasome machinery rather than cell type-specific factors.

These results validate Ctr-AIDE as a genome-integrated AID2 platform that enables precise, conditional depletion of *Chlamydia* effectors across diverse host cell lines while preserving native expression dynamics. By eliminating plasmid dependencies and leveraging the host ubiquitin-proteasome pathways, Ctr-AIDE establishes a new framework for dissecting essential bacterial virulence factors during infection.

### Ctr-AIDE Functions in Primary Cells

To validate the broad applicability of Ctr-AID, we established the Ctr-AIDE system in primary cells. Primary Murine Reproductive Tract (PMRT) organoids, derived from whole female reproductive tract digests, were transduced with lentivirus encoding OsTIR1(F74G) or OsTIR1(E7K/E10K/F74G) and selected for stable expression (Fig. S6). Cells derived from these organoids were then grown as 2D monolayers to facilitate 5-Ph-IAA treatment when used for Ctr-AIDE degradation assays. In PMRT cells expressing OsTIR1(F74G), we observed rapid, 5-Ph-IAA-dependent depletion of mAID-tagged Cdu1 comparable to cancer cell lines: immunoblotting confirmed near-complete Cdu1 loss within 1 hour of 5-Ph-IAA treatment, while controls (WT primary cells expressing dysfunctional OsTIR1(E7K/E10K/F74G), DMSO treated cells, and non-degron Cdu1 constructs) showed no degradation (Fig. 3A-B). Immunofluorescence microscopy further demonstrated auxin-driven disappearance of inclusion membrane-localized Cdu1-mAID and its restoration within 2 hours after 5-Ph-IAA washout (Fig. 3C). These results confirm that Ctr-AIDE operates robustly across both cancer and primary cell models, further prove its versatility for studying *Chlamydia* effector dynamics in physiologically relevant environments.

**Figure 3.**
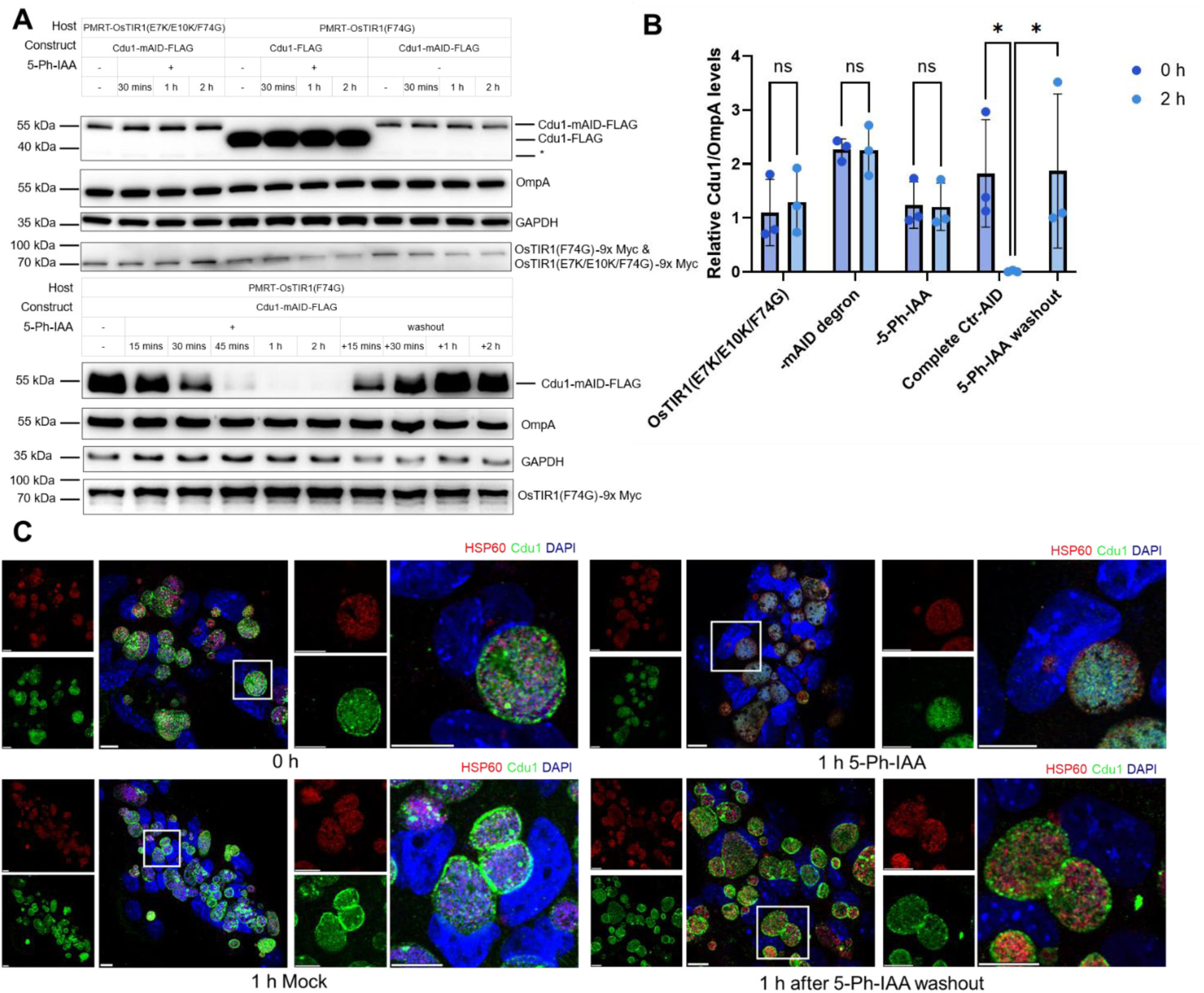
Ctr-AIDE can be applied in primary cells. (A) Immunoblot of the Ctr-AID-mediated regulation of Cdu1 levels in PMRT cells. Degradation treatment as in Figure 2, but a higher infection load (MOI = 10). Cdu1 and OsTIR1 mutants were detected with anti-FLAG and anti-Myc antibodies; OmpA and GAPDH served as loading controls. (B) Quantification of Cdu1 degradation in primary cells. FLAG levels (normalized to OmpA, mean ± SD) from three independent replicates (one shown in A). Significance assessed by two-tailed paired t-test (*p < 0.05; n.s., not significant). (C) Fluorescence microscopy of Ctr-AIDE dynamics in primary cells. Cdu1 (green, anti-FLAG), inclusions (Hsp60, red), and DNA (DAPI, blue). Scale bar = 10 µm. All experiments were replicated ≥3 times with consistent results.

### Assessing Cdu1 Function Dynamics

Leveraging the Ctr-AIDE system, we first investigated the deubiquitinase function of Cdu1, previously implicated in countering autophagy signaling ^35, 36^. We focused on the autophagy marker p62 to track inclusion membrane integrity and bypass potential confounding effects of Ctr-AIDE-induced K48-linked ubiquitination (Fig. S3). Depletion of Cdu1 triggered delayed recruitment of autophagy marker p62 to inclusions: the p62 signal on inclusions was minimal 1 hour after 5-Ph-IAA treatment but markedly increased by 4 hours (Fig. 4A). This phenotype was rapidly reversible with p62 recruitment dissipating within 1 hour of 5-Ph-IAA washout (Fig. 4A). Quantification of p62-positive inclusions confirmed this phenotype: Cdu1 depletion triggered p62 accumulation, while re-expression restored autophagy signaling evasion (Fig. 4B).

**Figure 4.**
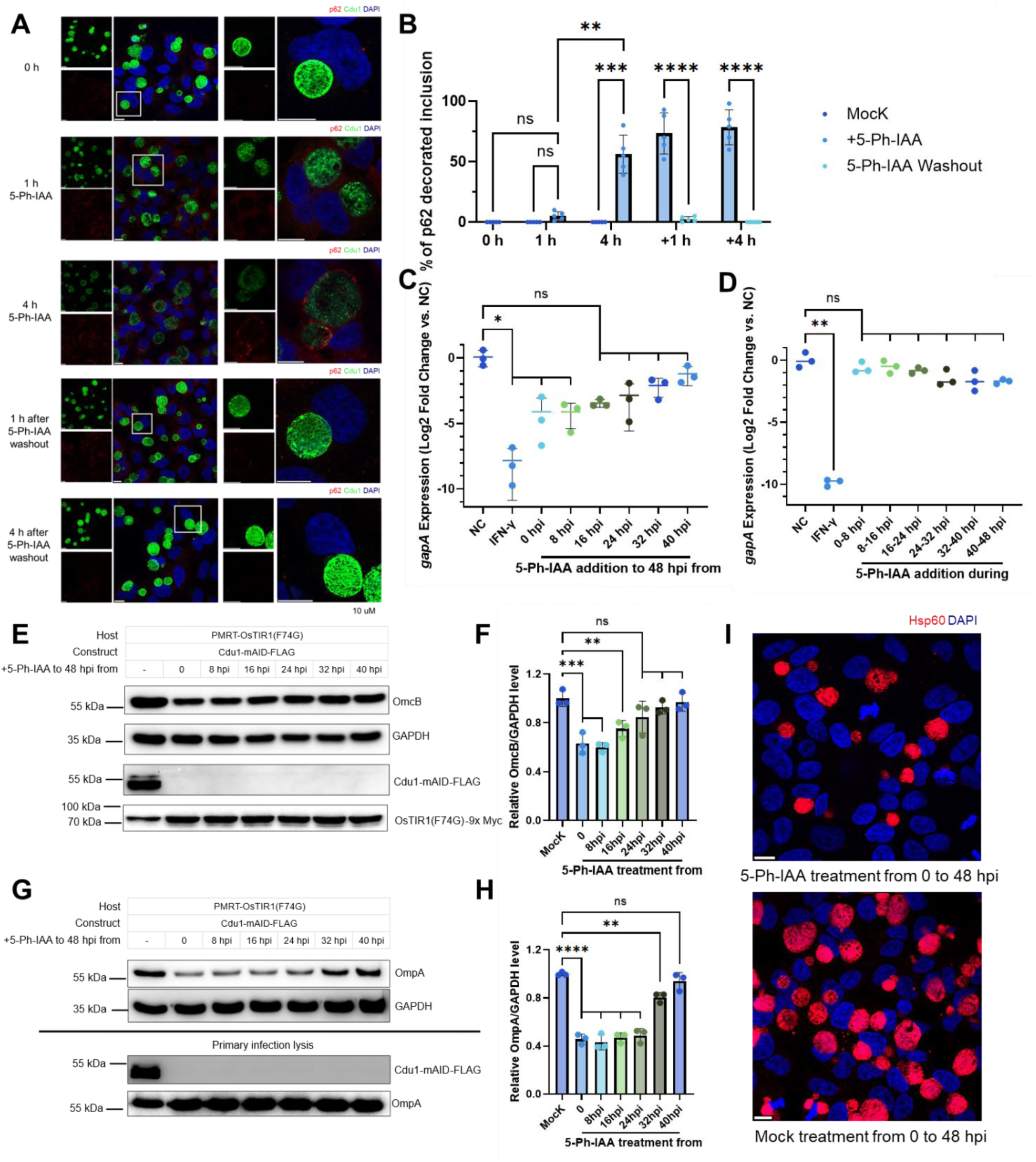
Cdu1 loss induces transient p62 accumulation and impairs *Ctr* metabolic activity and infectivity. (A) Spatiotemporal dynamics of p62 recruitment to *Ctr* inclusions upon Cdu1 depletion. Fluorescence microscopy of HeLa cells infected with Cdu1-mAID *Ctr* shows p62 (red, anti-SQSTM1) accumulation on inclusions after 4 h of 5-Ph-IAA treatment, dissipating within 1 h of Cdu1 re-expression. Cdu1 (green, anti-FLAG) and DNA (DAPI, blue) mark bacterial and host nuclei. Scale bar = 10 µm. (B) Quantification of p62-positive inclusions. Data represent mean percentage ± SD from 500 inclusions per condition across five independent experiments. Statistical significance was assessed by two-way ANOVA with Tukey’s test (****p < 0.0001, ***p < 0.001, **p < 0.01; n.s., not significant) comparing Cdu1 depletion/re-expression timepoints. +1 h and +4 h indicate hours after 5-Ph-IAA washout following 4 h treatment. (C) Cdu1 depletion reduces *Ctr* metabolic activity in primary cells. RT-qPCR of *gapA* (metabolic marker) and *rrs* (16S rRNA, control) at 48 hpi in Cdu1-mAID strains treated with 5-Ph-IAA at indicated timepoints. IFN-γ (50 U/mL) served as a positive control. Data normalized to DMSO-treated controls. Statistical analysis by one-way ANOVA with Dunn’s test (*p < 0.05; n.s.). (D) Short-term (8 h) depletion shows no effect (**p < 0.01; n.s.). (E) Cdu1 depletion before 48 hpi disrupts RB-to-EB redifferentiation. Immunoblot of OmcB (EB marker) in Cdu1-mAID strain degraded from indicated timepoints to 48 hpi. Cdu1 (anti-FLAG) and OsTIR1(F74G) (anti-Myc) confirm degradation; GAPDH serves as loading control. (F) Quantification of OmcB normalized to GAPDH (mean ± SD) from three replicates (one shown in E). Statistical significance assessed by one-way ANOVA with Dunnett’s test (***p < 0.001, **p < 0.01; n.s.). (G) Cdu1 loss reduces infectious progeny. Lysates from primary infection (MOI = 1, Cdu1 degraded from indicated timepoints to 48 hpi) were normalized by OmpA levels (lower “Primary infection lysis”), then used to infect HeLa cells. (H) Quantification shows progeny infectivity correlates with Cdu1 activity. OmpA normalized to GAPDH (mean ± SD, n = 3; one replicate shown in G). Statistical analysis by one-way ANOVA with Dunnett’s test (****p < 0.0001, **p < 0.01; n.s.). (I) Immunofluorescence microscopy confirms reduced progeny formation. HeLa cells infected with progeny from Cdu1-depleted (5-Ph-IAA 0–48 hpi) or DMSO-treated *Ctr* were imaged at 24 hpi. Inclusions (Hsp60, red) and DNA (DAPI, blue). Scale bar = 10 µm. All experiments were replicated ≥3 times with consistent results.

Previous studies reported that Cdu1 dysfunction impairs chlamydial growth in primary cells, but not in HeLa cells (Fig. S7) ^36^. To evaluate Cdu1’s role in bacterial growth, we implemented two degradation strategies in the 2D PMRT cell model: (i) continuous degradation, in which 5-Ph-IAA treatment was initiated at 0, 8, 16, 24, 32, or 40 hpi and maintained until 48 hpi, and (ii) acute 8-hour pulses, initiated at staggered 8-hour intervals, and lasting 8 hours each (Fig. S8). Metabolic activity was monitored via RT-qPCR at 48 hpi for *gapA* (encoding glyceraldehyde-3-phosphate dehydrogenase) ^44^. Sustained depletion significantly reduced *gapA* expression when initiated before 16 hpi (Fig. 4C). In contrast, short-term 8-hour degradation had no significant effect, indicating that Cdu1‘s metabolic role is only compromised by prolonged depletion (Fig. 4D).

Cdu1 was also required for *Ctr* developmental progression ^36^. During infection, bacteria transition from the infectious, non-replicating elementary body (EB) to the replicative, non-infectious reticulate body (RB). Depletion before 24 hpi reduced levels of OmcB, a late-stage EB marker (Fig. 4E-F), coinciding with the onset of RB-to-EB differentiation ^45^. These findings highlight Cdu1’s critical role in initiating developmental transitions.

To assess the consequence of reduced EB transitions, we also quantified progeny formation by performing infectivity assays (Fig. S9). Chlamydial lysates from primary cell infections were used to infect fresh HeLa cells. Progeny derived from cultures in which Cdu1 was depleted no later than 32 hpi exhibited significantly reduced OmpA levels in fresh cells at 24 hpi compared to untreated controls (Fig. 4G-I; p < 0.05), indicating that Cdu1 loss disrupts RB redifferentiation into infectious EBs. Interestingly, no reduction in progeny formation was observed following acute 8-hour degradation pulses (Fig. S10), reinforcing the requirement of sustained Cdu1 activity to support productive infection.

### Induced degradation and re-expression of IncA

Previous studies demonstrated that the C-terminus of IncA mediates inclusion infusion ^38^. To investigate the dynamics of IncA function as a key regulator of inclusion fusion and stability ^38, 39^, we applied the AIDE platform to this Inc protein. Using the same genome-integrated tagging strategy previously developed for Cdu1 (Fig. S11), we generated *Chlamydia* strains expressing FLAG-tagged IncA or IncA-mAID-FLAG (Fig. S1B).

Addition of 5-Ph-IAA induced rapid degradation of IncA-mAID within 1 hour, followed by re-expression within 30 minutes after 5-Ph-IAA washout. In addition, degradation required a functional E3 ligase OsTIR1(F74G), the mAID degron and addition of 5-Ph-IAA, since controls lacking any component or with dysfunctional E3 ligase retained stable IncA levels (Fig. 5A-B). Immunofluorescence microscopy also demonstrated the loss of inclusion membrane-localized IncA 1 hour after addition of 5-Ph-IAA and its restoration after 5-Ph-IAA washout (Fig. 5C).

**Figure 5.**
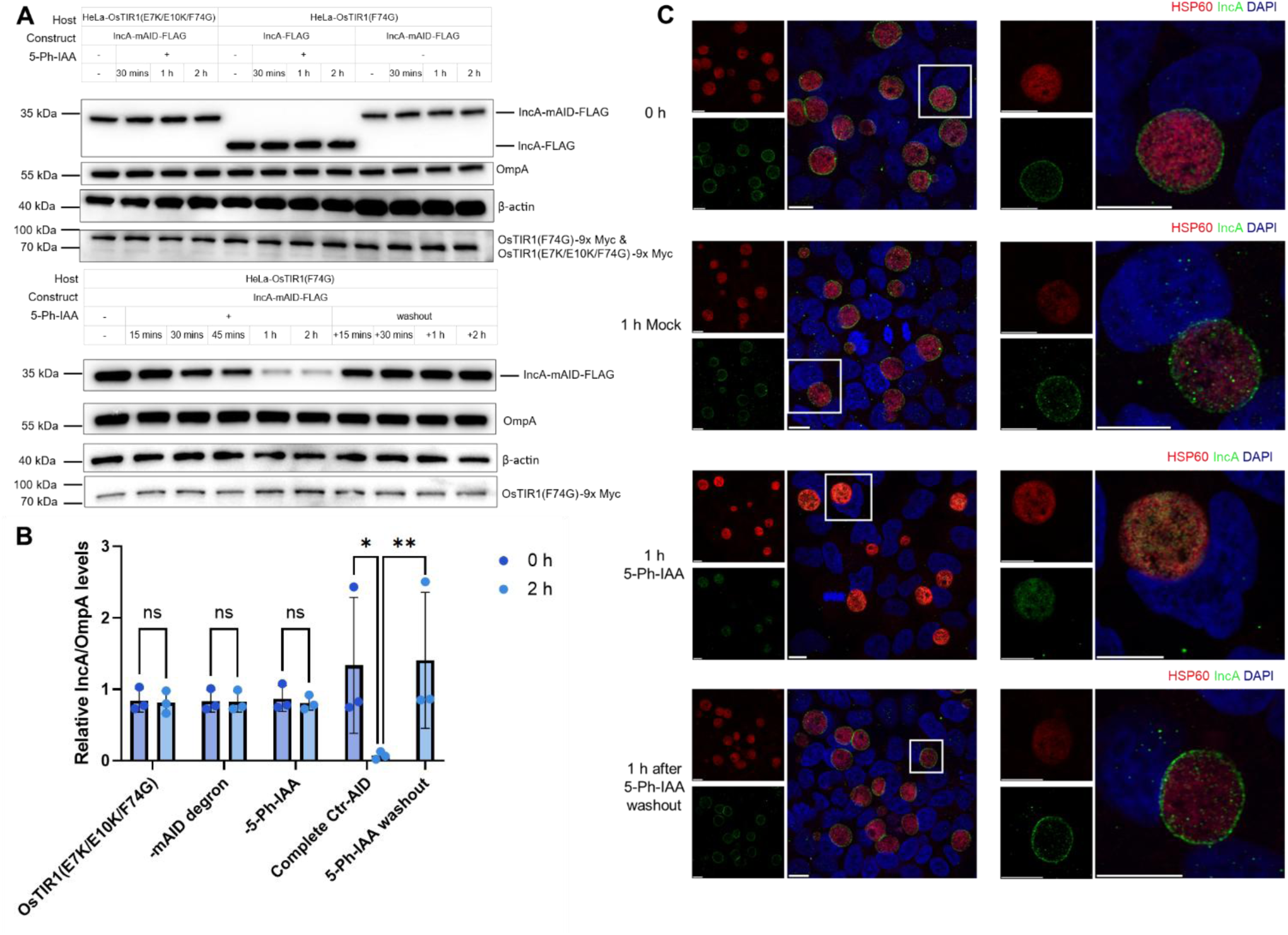
Ctr-AIDE system enables conditional control of IncA expression in *Chlamydia*. (A) Ctr-AID-mediated regulation of IncA-mAID expression. Degradation and re-expression protocols were performed as described for Cdu1. IncA (anti-FLAG) and OsTIR1 mutants (anti-Myc) levels were monitored, with OmpA as a loading control and β-actin as a host cell control. (B) Quantification of IncA degradation. FLAG levels normalized to OmpA (mean ± SD) from three independent immunoblot replicates (one shown in A). Significance assessed by two-tailed paired t-test (****p < 0.0001; n.s., not significant). (C) Immunofluorescence microscopy of IncA depletion dynamics. IncA (green, anti-FLAG), inclusions (Hsp60, red), and DNA (DAPI, blue). Scale bar = 10 µm. All experiments were repeated ≥3 times with consistent results.

### Accessing IncA Function Dynamics

We next evaluated the functional consequences of IncA depletion. Consistent with previous reports ^38, 39, 46^, IncA degradation by addition of 5-Ph-IAA from 0-24 hpi abolished inclusion fusion, yielding multiple or fragmented inclusions (Fig. 6A). This confirms IncA’s essential role in mediating homotypic inclusion fusion.

**Figure 6.**
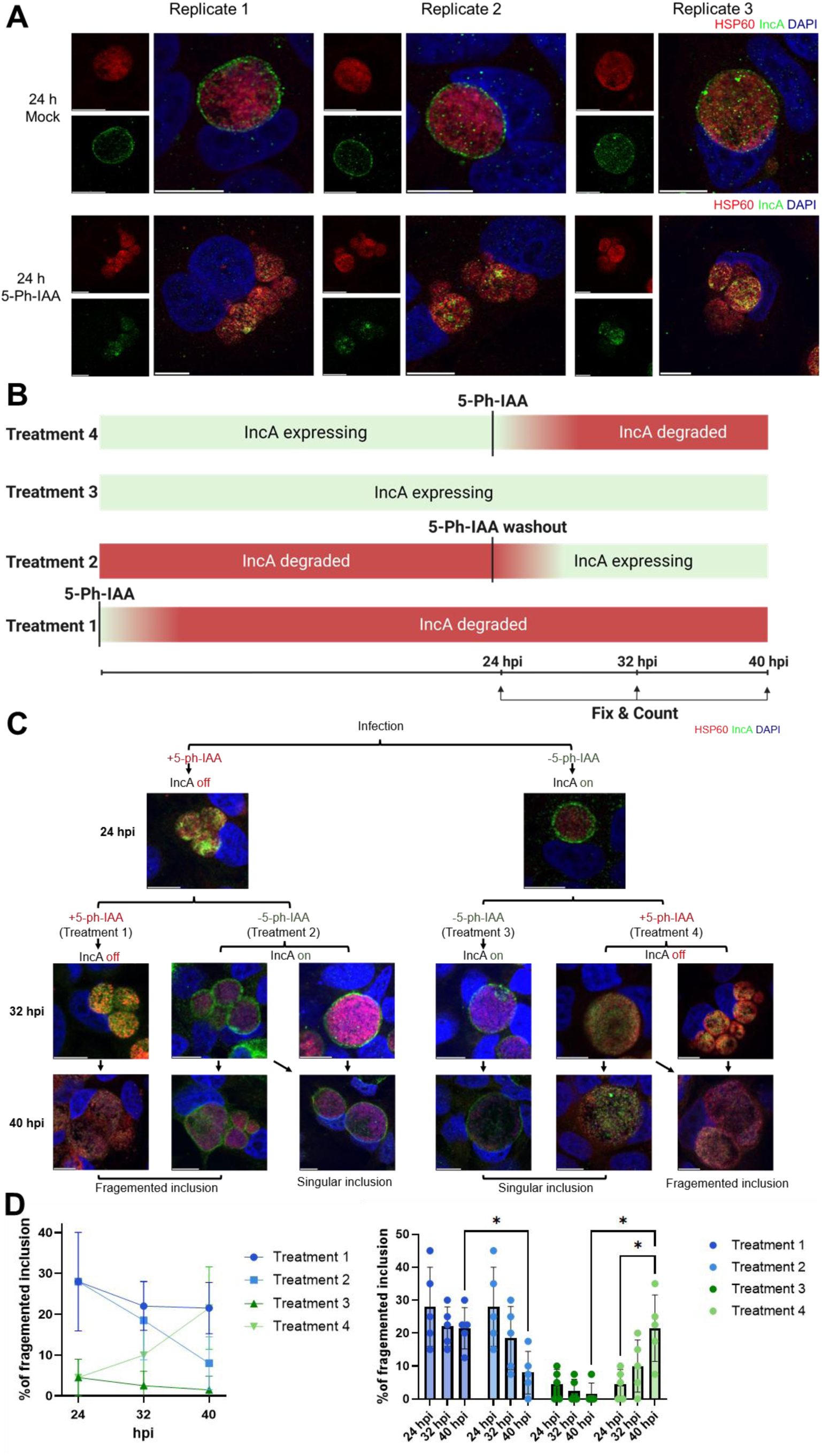
IncA degradation reveals its essential role in inclusion fusion during the *C. trachomatis* life cycle. (A) IncA degradation prevents inclusion fusion. HeLa cells infected with IncA-mAID strains were treated with 5-Ph-IAA (IncA degraded) or DMSO (control) from infection to 24 hpi. Degraded samples show fragmented inclusions, whereas controls form single fused inclusions. IncA (green, anti-FLAG), inclusions (Hsp60, red), and DNA (DAPI, blue). Scale bar = 10 µm. (B) Schematic of IncA termination and re-expression experiments. (C) Immunofluorescence microscopy of inclusion morphology under different life cycle time points under Treatment 1-4. (D) Quantification of host cells containing multiple intact inclusions (≥2) upon Treatment 1-4. Data represents mean percentage ± SD from 200 cells counted per condition across five independent biological replicates. Significance assessed by two-way ANOVA with Tukey Multiple comparisons test (*p < 0.05; not significant was not shown). All experiments were replicated ≥3 times with consistent results.

Despite intensive research on the mechanism of inclusion fusion ^39, 46, 47^, it remains unclear whether IncA activity is continuously required to maintain fused inclusions or becomes dispensable once fusion is complete. Addressing this question has been challenging due to the long stability of membrane-integrated IncA, which limits the effectiveness of conventional silencing approaches. Continuous sRNA-mediated silencing of IncA expression from 0–24 hpi effectively reduced IncA levels, whereas silencing initiated at 24 hpi failed to deplete the protein, which remained stable for at least another 24 h under these conditions (Fig. S12A-B). In contrast, AIDE enabled rapid depletion of membrane-integrated IncA within 1 h at 24 hpi (Fig. S12C-D), providing a powerful tool to directly assess the temporal requirements of IncA function. To determine whether IncA’s function is static or dynamic, we implemented time-resolved degradation schemes. When IncA was expressed from 0-24 hpi and then degraded from 24-40 hpi, inclusions fragmented into multiple smaller inclusions (Fig. 6C, Treatment 4), indicating that ongoing IncA activity is required to maintain inclusion integrity even after initial fusion. To further dissect these temporal dependencies, we compared two conditions: (i) Expression pause: IncA expressed from 0-24 hpi, then degraded until 40 hpi (Treatment 4). (ii) Rescue: IncA degraded from 0-24 hpi, then re-expressed via 5-Ph-IAA washout until 40 hpi (Treatment 2). Controls included continuous IncA expression (0-40 hpi, Treatment 3) or degradation (0-40 hpi, Treatment 1) (Fig. 6B).

Quantitative analysis revealed progressive inclusion fragmentation in the ‘Expression pause’ group (Treatment 4) after 5-Ph-IAA addition at 24 hpi, while the ‘Rescue’ group (Treatment 2) restored inclusion fusion after 5-Ph-IAA washout at 24 hpi. Control treatments confirmed that continuous IncA expression maintained fused inclusions (Treatment 3), whereas sustained IncA degradation (Treatment 1) led to irreversible fragmention (Fig. 6C-D). Together, these results demonstrate that IncA is continuously required to preserve inclusion integrity, acting persistently to counteract host-driven membrane destabilization.

## Discussion

Traditional genetic tools (e.g., classical knockout, transposon mutagenesis, silencing technologies) have proven inadequate for resolving the dynamic function of secreted proteins in pathogenic bacteria ^48–52^. Conditional tools such as CRISPRi and engineered sRNAs allow inducible repression but are hindered by issues including incomplete knockdown, promoter leakiness, plasmid stability and expression constraints, off-target and polar effects, delayed phenotypes for stable proteins, and challenges in restoring target gene expression ^31, 49, 53^. By repurposing the host ubiquitin-proteasome system, AIDE overcomes these limits, enabling rapid (minutes-wise), reversible depletion of bacterial effectors at the host-pathogen interface. To our knowledge, this work represents the first successful application of TPD to host-delivered bacterial effectors, addressing a decades-old methodological gap.

One of the most notable advantages of AIDE lies in its versatile application which circumvents the complex molecular design required to establish functional PROTACs for targeted protein degradation, which is a prohibitive barrier for studying many secreted effector proteins whose structures remain uncharacterized ^54, 55^. Its minimalist design, featuring a small 7 kDa mAID degron fused to targets that need to be accessible to host cells cytosol, ensures seamless genomic integration and preserves native protein function, as demonstrated for Cdu1 and IncA. AIDE also bypasses several shortcomings of bacteria-specific strategies like BacPROTACs, which depend on poorly defined bacterial proteases (e.g., ClpCP) ^56^, and the delivery of compounds over two or more membranes. Unlike transcriptional or translational knockdown methods, AIDE operates post-translationally, avoiding off-target effects on adjacent genes ^57, 58^.

AIDE uniquely enables causal analysis of secreted effector function at defined times, a capability lacking in conventional transcriptional or translational silencing methods such as CRISPRi or engineered sRNAs. Although these approaches can suppress expression when initiated early and allow re-expression after inducer washout, they rarely achieve rapid depletion of pre-synthesized effectors during the mid to late infection cycle (Fig. S12). This limitation arises from the long half-lives of membrane-associated proteins and pathogen-encoded countermeasures such as the deubiquitinase Cdu1. As a result, two constraints emerge: (i) phenotypes are typically revealed only upon re-expression rather than acute loss, and (ii) prolonged early inhibition can drive bacterial adaptation and persistence, confounding interpretation of single-effector loss ^59^. By contrast, AIDE affords minute-scale, reversible control, enabling immediate degradation and rapid re-expression across precisely defined infection windows, with the potential for repeated on/off toggling. These properties reveal effector dynamics previously inaccessible to silencing-based strategies, as demonstrated for Cdu1 and IncA.

We demonstrate that the ubiquitin-directed degradation of transmembrane effectors harnesses the host p97 (VCP) system to enable efficient, proteasome-dependent clearance. p97 facilitates this process by actively extracting ubiquitinated substrates from membranes and delivering them to the cytosolic proteasome for degradation ^43^. This mechanism likely confers exceptional efficiency in depleting otherwise stable, membrane-anchored proteins, an effect not achievable with conventional knockdown approaches. Consistent with this, we detected Cdu1-mAID ubiquitination as early as 5 minutes after 5-Ph-IAA addition, with protein levels reduced by ∼70% just 15 minutes later (Fig. 1D). Similarly, Ctr-AIDE enabled near-complete depletion of the stably membrane-integrated effector IncA within 1 hour (Fig. 5). Auxin washout allows cytoplasmic proteins to rapidly fold and be re-exported by the T3SS, restoring membrane activity for near-instantaneous functional analysis, as demonstrated by Cdu1’s suppression of autophagy signaling within 1 hour. Importantly, Ctr-AIDE induced rapid, minute-scale ubiquitin conjugation and degradation even of the deubiquitinase Cdu1. This finding was unexpected, as (i) the auto-acetyltransferase activity of Cdu1 on lysine residues has been proposed to protect it from K48-linked ubiquitination ^37^, and (ii) its potent, broad K48-directed deubiquitinase activity would be expected to counteract K48-mediated proteasomal degradation.

By applying Ctr-AIDE to *Chlamydia* Incs, we resolved the spatiotemporal dynamics of two critical effectors. Cdu1 functions as a dual-phase sentinel, coordinating its enzymatic activities to safeguard the inclusion niche. Intriguingly, autophagy marker accumulation was delayed until 4 hours post-Cdu1 degradation, with no significant increase observed within the first hour. We attribute this delay to Cdu1’s acetyltransferase activity. Previous studies indicate that Cdu1-mediated acetylation protects distinctive Incs from host ubiquitination ^37, 60^. Consequently, during the initial hour post-degradation, these effectors remained acetylated and resistant to ubiquitination. Over time, however, host deacetylases (e.g., HDACs) progressively deacetylated Incs, rendering Incs susceptible to ubiquitin-mediated tagging ^61, 62^. In contrast, re-expression experiments revealed that only mature, inclusion membrane-localized Cdu1 clears autophagy markers within 1 hour, demonstrating its rapid, compartment-specific activity. This highlights its role as a frontline defender against host immune tagging.

During mid-late development (16-48 hpi), Cdu1 transitions to sustaining nutrient acquisition and bacterial growth and development. Time-resolved degradation showed that Cdu1 is essential in this window, coinciding with inclusion expansion, nutrient scavenging, and RB-to-EB differentiation. Because only inclusion membrane-localized Cdu1 is degraded, and previous studies show that growth defects in Cdu1-deficient strains are not caused by autophagy-mediated clearance, we propose that Cdu1 degradation during this critical window impairs Golgi vesicle recruitment ^36^. This, in turn, starves replicating RBs of lipids, nucleotides, and ATP required both for binary fission and for the energy-intensive conversion into infectious EBs, ultimately reducing progeny yield and infectivity. In summary, the Cdu1’s instant autophagy clearance and sustained nutrient acquisition emphasize its flexibility and its important role in *Ctr* developmental progression.

Conversely, IncA maintains inclusion architecture through persistent activity of its SNARE-like domain (SLD1/2). These domains competitively engage host SNARE proteins, blocking lysosomal fusion while driving homotypic vacuole mergers. Strikingly, following inclusion fusion, degradation of inclusion membrane-bound IncA triggered vacuole fragmentation, a phenotype reversed by IncA re-expression. This dynamic equilibrium demonstrates that inclusion stability requires continuous IncA activity to counteract host membrane destabilization, not only initial fusion.

These insights exemplify how Ctr-AIDE bridges temporal resolution with subcellular precision, unmasking effector roles in developmental timing and niche stability. The system’s capacity to decouple sequential functions such as autophagy evasion, nutrient scavenging, and membrane stabilization reveals stage-specific therapeutic vulnerabilities. Targeting these phases could halt *Chlamydia’s* developmental cycle, offering a roadmap for precision antimicrobial design.

More broadly, AIDE harnesses eukaryotic machinery to control pathogen proteins, a host-directed strategy that opens new avenues for manipulating intracellular bacteria. While sophisticated protein-control systems are well established in human cells, they are difficult to implement in bacteria, particularly intracellular species with limited genetic tractability ^63^. By contrast, host-directed strategies such as AIDE directly exploit host-encoded pathways to regulate pathogen proteins with temporal precision, independent of the pathogen’s biology, and can potentially circumvent microbial defense systems.

At present, AIDE targets inclusion membrane proteins that are directly accessible from the host cytosol. In combination with inducible delivery systems, analogous approaches could be extended to effectors located within the bacterial cytosol ^64–66^.

Conceptually, AIDE also shifts the balance of control from the pathogen to the host. Pathogenic infections are notoriously difficult to cure because evolved microbes co-opt host pathways for their own benefit. Tools such as AIDE establish a synthetic defensive system that reverses this hierarchy, enabling host cells to dictate pathogen function. With continued advances in targeted protein degradation, it may soon become possible to regulate pathogen proteins without engineered degrons, by instead recruiting specific host E3 ligases. This trajectory offers a promising framework for the development of next-generation anti-pathogen therapeutics.

In conclusion, AIDE redefines the mechanistic dissection of host-pathogen interactions by merging precision degradation with temporal control. Its success in revealing the dynamic functions of *Chlamydia* effectors highlights the immense potential of host-directed tools to accelerate antimicrobial discovery targeting pathogen-secreted effector proteins. Ctr-AIDE therefore establishes a blueprint for studying effector dynamics in the very large group of pathogenic bacteria that secret effector proteins into host cells.

## Methods

### Antibodies used

Antibodies and their working dilution are listed in Table S5.

### Plasmids and *Escherichia coli* strains

All plasmids and primers are listed in Supplementary Tables S2 and S4. Plasmid construction proceeded as follows: The *Cdu1* coding sequence (*CTL0247*) and its 250 bp promoter region were amplified from *Ctr* serovar L2/434/Bu genomic DNA. mAID degron and FLAG-tag sequences were fused to the C-terminus of Cdu1 via PCR using primers with 30 bp homology overlaps. For genomic knock-in, the pKW-L2 backbone ^40^ was modified by Gibson Assembly (New England Biolabs) to incorporate: (i) 3 kb homology arms flanking Cdu1, (ii) engineered Cdu1-mAID-FLAG or Cdu1-FLAG inserts, (iii) aadA-GFP selection cassette (spectinomycin resistance/GFP fusion).

Identical strategies were applied for IncA (*CTL0374*) knock-in constructs, with all assemblies verified by Sanger sequencing.

*Escherichia coli* strains (Supplementary Table S3) were cultured in Lysogeny Broth (LB; Carl Roth) or on LB agar plates (1.5% w/v agar) at 37°C. DH10B served as the host for plasmid propagation. Media were supplemented with ampicillin (100 μg/mL) or spectinomycin (50 μg/mL) as required. Glycerol stocks (50% v/v) of engineered strains were stored at −80°C.

### Culture of organoids and organoid-derived 2D monolayers

3 adult (7-8 weeks old) female C57BL/6J mice were euthanized under institutional ethical approval. The entire reproductive tract (ovaries, oviducts, uterus and cervix) was carefully excised, washed twice in ice-cold PBS containing 1% penicillin/streptomycin, and any excess adipose or connective tissue was trimmed away.

Tissues were minced into approximately 1–2 mm pieces and digested in digestion buffer (PBS + 1 mg/mL collagenase I [Thermo Fisher] + Trypsin 5% [Gibco] + 50 µg/mL DNase I [New England Biolabs]) at 37 °C for 60 minutes with gentle rocking. After digestion, the cell/tissue slurry was passed through a 40 µm nylon strainer, washed once in PBS. Cells were collected by centrifugation (1,000 × g, 5 min, 4°C), then resuspended in Matrigel. Aliquots of 50 µL Matrigel domes were plated into pre-warmed 24-well plates and each well received 500 µL of organoid medium (prepared as described in ^67^). Cultures were maintained at 37°C under 5% CO₂ in a humidified incubator.

Organoids were passaged every 7-14 days at splitting ratios ranging from 1:2 to 1:10, based on growth density, with medium replenished every 2-3 days. For organoid passage, briefly, Matrigel domes were dissolved in ice-cold organoid medium. Released organoids were collected by centrifugation (400 × g, 5 min, 4°C) and digested in 1 mL of TrypLE Express (Thermo Fisher Scientific) for 15 minutes at 37°C with intermittent pipette trituration to obtain small clusters. Digested cells were pelleted again, resuspended in Matrigel, and plated in 24-well plates. Finally, 500 µL of fresh organoid medium was added to each well.

To generate 2D monolayers, organoids were released from Matrigel and dissociated into single cells using TrypLE at 37°C for 10 minutes. Dissociated cells were seeded into CellAdhere™ Type I Collagen-pretreated microwell plates or flasks (STEMCELL Technologies) and maintained in organoid medium under identical conditions.

### Cancer cell lines and cultivation

All the cell lines are listed in Supplementary Tables S1. HeLa 229, A-375, HCT 116, U-2 OS, and McCoy cells were maintained in RPMI-1640 medium (Gibco) supplemented with 10% fetal calf serum (FCS; Paa Laboratories), while HEK293T cells were cultured in DMEM (Dulbecco’s Modified Eagle Medium; Gibco) with 10% FCS. PMRT cells were grown in custom organoid medium as previously described. All cells were incubated at 37°C under 5% CO₂ humidified conditions.

### Lentivirus generation and transduction

Lentivirus constructs for stable OsTIR1(F74G) or OsTIR1(E7K/E10K/F74G) expression were produced in HEK293T cells co-transfected with pRRL_OsTIR1F74G or pRRL_OsTIR1(E7K/E10K/F74G), and helper plasmids psPAX2/pMD2.G using Lipofectamine 3000 (Thermo Fisher). Cancer cell lines were transduced with viral supernatants containing 5 μg/mL Polybrene (Merck Millipore). Organoids were transduced as previously published ^68^ with the following modifications: organoid medium formulation was maintained as described above, while Chir99021 and Y27632 were omitted. Transduced organoids underwent spinoculation (500 × g, 37°C, 60 min) followed by 3-hour static incubation at 37°C.

Following two infection rounds, hygromycin selection (1 mg/mL; Carl Roth) was applied to all cells (cancer and organoid) in sequential 24-hour PBS wash and medium renew cycles to eliminate untransduced cells.

### *Ctr* propagation and transformation

*Ctr* serovar L2 (434/Bu) was propagated in McCoy cells. For infections, host cells at 70% confluency were incubated with EBs at an MOI of 1 in RPMI-1640/10% FCS at 35°C under 5% CO₂. Bacterial stocks were prepared by harvesting infected cells at 48 hpi via mechanical disruption with glass beads. Debris was pelleted by centrifugation (1,000 × g, 10 min, 4°C), and EBs were subsequently isolated from supernatants by ultracentrifugation (30,000 × g, 30 min, 4°C). Purified EBs were resuspended in sucrose-phosphate-glutamic acid (SPG) buffer (75 g/L sucrose, 0.52 g/L KH₂PO₄, 1.22 g/L Na₂HPO₄, 0.72 g/L L-glutamic acid, pH 7.4) and stored at −80°C.

Transformation was performed as described ^69^. Briefly, 10 µg plasmid DNA and 1.6 × 10⁷ inclusion-forming units (IFUs) of EBs were mixed in CaCl₂ buffer (10 mM Tris-HCl pH 7.4, 50 mM CaCl₂), then incubated with 8 × 10⁶ McCoy cells for 20 minutes at room temperature. The mixture was transferred to T75 flasks containing McCoy’s 5A medium (Gibco) with 10% FCS. At 48 hpi, lysates were passaged onto fresh McCoy cells under selection with 500 µg/mL spectinomycin and 1 µg/mL cycloheximide (both Carl Roth), with passages repeated every 48 hours. Recombinant strains were verified by fluorescence and immunoblotting (Supplementary Table S3).

### Degradation assays

OsTIR1(F74G)- or OsTIR1(E7K/E10K/F74G)-expressing cancer cell lines were seeded in 12-well plates and infected with specified *Chlamydia* strains at MOI = 1. Degradation was induced by adding 1 µM 5-Ph-IAA (5-phenylindole-3-acetic acid; MedChemExpress), 200 ng/mL anhydrotetracycline (aTC; Merck Millipore) or DMSO control to the culture medium. To terminate degradation, cells were washed once with DPBS and replenished with fresh drug-free medium. For primary cell lines, identical protocols were applied except for higher infection loads (MOI = 10).

All samples including Mock (untreated controls) were harvested at 28 hpi for further analysis, with degradation and washout timed to synchronize endpoints (Fig. 2A). For multi-phase degradation-recovery experiments, treatments were applied in defined windows prior to harvest.

### Infectivity and metabolic assays

HeLa cells and primary PMRT cells were seeded in 12-well plates and infected with specified *Ctr* strains at MOI = 1. For metabolic assessment at 48 hpi, cells were lysed with 0.5 mm sterile glass beads in fresh HBSS; lysates were centrifuged (1,000 × g, 10 min, 4°C) and supernatants aliquoted for RT-qPCR (*gapA* quantification) and OmcB immunoblotting. For progeny infectivity assays, parallel 48 hpi lysates were normalized by OmpA immunoblotting to equalize bacterial loads by adding fresh HBSS. Normalized supernatants were used to infect fresh HeLa monolayers, with progeny harvested at 24 hpi for immunoblot analysis.

### Inhibitor and IFN-γ treatments

Proteasome inhibitors (MG-132 at 10 µM, bortezomib at 1 µM; Cell Signaling), p97 inhibitor CB-5083 (10 µM; MedChemExpress), and ubiquitin-activating enzyme inhibitor TAK-243 (1 µM; TAK-243) were prepared as 1,000× stock solutions in DMSO. Inhibitors were diluted to working concentrations in culture medium and applied 3 hours prior to 5-Ph-IAA treatment, and kept in medium during the 1 hour 5-Ph-IAA treatment. For IFN-γ induction, cells were pretreated with 50 U/mL recombinant human IFN-γ (Merck Millipore) for 2 hours before infection.

### Indirect immunofluorescence

Immunofluorescence was performed on cells grown on glass coverslips. Following treatments or infections, cells were fixed with 4% paraformaldehyde (PFA; Carl Roth) for 15 minutes at room temperature, permeabilized with 0.2% Triton X-100 in PBS (30 min), and blocked in PBS containing 2% FCS (1 hr). Primary antibodies diluted in blocking buffer were applied for 1 hour at room temperature. After three 5-minutes PBS washes, secondary antibodies were incubated for 1 hour protected from light. Coverslips were mounted using Fluoroshield mounting medium (Abcam) and imaged on a Leica STELLARIS 5 confocal microscope (63×/1.4 NA oil objective, inverted configuration). Images were processed using Leica *LAS X* software (3.10.0).

### Post-immunostaining gel embedding and expansion

Immunostained samples were post-fixed in 0.25% glutaraldehyde (10 mins; Carl Roth). Specimens were embedded in monomer solution (1.375 M sodium acrylate, 2.5% acrylamide, 0.15% bis-acrylamide, 2 M NaCl, 1× PBS pH 7.4, 0.2% APS, 0.2% TEMED; all Sigma-Aldrich) and polymerized (1 hr). Gels were digested in proteinase K buffer (8 U/mL in 50 mM Tris pH 8.0, 1 mM EDTA, 0.5% Triton X-100, 0.8 M guanidine HCl; 37°C, overnight), then expanded in deionized water until saturation (4-5× linear expansion). Expanded gels were imaged using a Leica STELLARIS 5 confocal microscope (63×/1.4 NA oil objective, inverted configuration). Images were processed using Leica *LAS X* software (3.10.0).

### SDS-PAGE and immunoblotting

Cell lysates were prepared in RIPA Lysis and Extraction Buffer (Thermo Fisher Scientific) supplemented with Protease and Phosphatase Inhibitor Cocktail (Thermo Fisher Scientific). Lysates were clarified by centrifugation (16,000 × g, 10 min, 4°C), and supernatants were mixed with ROTI®Load 1 loading buffer (Carl Roth) followed by denaturation at 94°C for 5 minutes. Proteins were resolved on 12% SDS-PAGE gels and electrophoretically transferred to nitrocellulose membranes using a Bio-Rad Trans-Blot Turbo system (7 V, 25 minutes). Membranes were blocked with EveryBlot Blocking Buffer (Bio-Rad) for 10 minutes at room temperature, followed by overnight incubation with HRP-conjugated primary antibodies diluted per manufacturer specifications. After three 5-minute washes with TBS-T (0.1% Tween-20), blots were developed with Immobilon Forte Western HRP Substrate (Merck Millipore) for 1 minute and imaged using an Intas Chem HR 16-3200 system (Auto exposures). Protein band intensities were quantified by densitometry using ImageJ (v1.53) with background subtraction.

### Quantitative Real-Time PCR (qRT-PCR)

Total RNA was extracted using the RNeasy Mini Kit (Qiagen). cDNA was synthesized from 1 μg RNA using the High-Capacity cDNA Reverse Transcription Kit (Thermo Fisher Scientific) in 20 μL reactions (25°C for 10 min, 37°C for 120 min, 85°C for 5 min). qPCR was performed with SYBR Green Master Mix (Applied Biosystems) on a StepOnePlus™ system (Applied Biosystems) under standard cycling conditions: 95°C for 10 minutes, followed by 40 cycles of 95°C for 15 seconds and 60°C for 1 minute. Primer sequences are in Supplementary Table S2. Relative expression was calculated by the 2−ΔΔCt method with chlamydial *rrs* (16S rRNA) as the endogenous control.

### Statistics and data

Statistical analyses were performed using GraphPad Prism v10.4.0 (GraphPad Software). Data represent SD from ≥3 independent biological replicates (specified in figure legends). Normality and homoscedasticity assumptions were verified for all parametric tests. Significant differences (α = 0.05) were determined by two-tailed Student’s t-test (two groups) or one/two-way ANOVA with Dunnett’s/Tukey’s post hoc tests (multiple comparisons), as appropriate for experimental designs. Exact p-value thresholds, statistical tests, and degrees of freedom are provided in the corresponding figure legends.

Figures were assembled in Microsoft PowerPoint using graphical outputs from GraphPad Prism; Biological schematics were created with Microsoft PowerPoint and BioRender.com.

Supplementary tables contain the following primary datasets:

Supplementary Table S6: qRT-PCR raw Ct values and curve data from ‘Design and Analysis’ program

Supplementary Table S7: Quantitative analysis of inclusion fragmentation during IncA degradation

Supplementary Table S8: p62-positive inclusion quantification during Cdu1 degradation

Supplementary Table S9: Immunoblot densitometry raw data

