## Supplemental information for "Minute-scale control of ubiquitin-mediated degradation reveals dynamics of bacterial secreted effector-functions"

Content:

Supplementary Table S1
Supplementary Table S2
Supplementary Table S3
Supplementary Table S4
Supplementary Table S5
Supplementary Figures S1-S11
Supplemental References

Table S1. Cell lines used in this study

| Cell line | Description | Source |
|---|---|---|
| McCoy | ATCC CRL-1696 | [1] |
| HeLa 229 | ATCC CCL-2.1 | [2] |
| HEK293T | ATCC CRL-3216 | [3] |
| A-375 | ATCC CRL-1619 | [4] |
| HCT 116 | ATCC CCL-247 | [5] |
| U-2 OS | ATCC HTB-96 | [6] |
| PMRT | Primary Murine Reproductive Tract cells | This study |
| HeLa-OsTIR1(F74G) | HeLa cells stably express OsTIR1(F74G)-9x Myc | This study |
| HeLa-OsTIR1(E7K/E10K/F74G) | HeLa cells stably express OsTIR1(E7K/E10K/F74G)-9x Myc | This study |
| A-375-OsTIR1(F74G) | A-375 cells stably express OsTIR1(F74G)-9x Myc | This study |
| HCT 116-OsTIR1(F74G) | HCT 116 cells stably express OsTIR1(F74G)-9x Myc | This study |
| U-2 OS-OsTIR1(F74G) | U-2 OS cells stably express OsTIR1(F74G)-9x Myc | This study |
| PMRT-OsTIR1(F74G) | Primary Murine Reproductive Tract cells stably express OsTIR1(F74G)-9x Myc | This study |
| PMRT-OsTIR1(E7K/E10K/F74G) | Primary Murine Reproductive Tract cells stably express OsTIR1(E7K/E10K/F74G)-9x Myc | This study |

Table S2. Plasmids used in this study

| Name | Description | Source |
|---|---|---|
| pKW-L2 | Suicidal plasmid for FRAEM genome-editing | [7] |
| pHL18 | pCMV-N1 plasmid expressing GFP-mAID-NLS-FLAG | This study |
| pHL20 | pCMV-N1 plasmid expressing GFP-NLS-FLAG | This study |
| pHL54 | pKW-L2-3kb upstream-Cdu1-mAID-FLAG-3kb downstream | This study |
| pHL55 | pKW-L2-3kb upstream-Cdu1-FLAG-3kb downstream | This study |
| pHL92 | pKW-L2-3kb upstream-IncA-mAID-FLAG-3kb downstream | This study |
| pHL93 | pKW-L2-3kb upstream-IncA- FLAG-3kb downstream | This study |
| pRRL_OsTIR1F74G | Plasmid used for producing Lentivirus expressing OsTIR1(F74G)-9x Myc | This study |
| pRRL_OsTIR1F74G_2mut | Plasmid used for producing Lentivirus expressing OsTIR1(E7K/E10K/F74G)-9x Myc | This study |
| psPAX2 | Lentivirus production helper plasmid | Addgene, Cat. # 12260 |
| pMD2.G | Lentivirus production helper plasmid | Addgene, Cat. # 12259 |

Table S3. Strains used in this study

| Strain | Description | Source |
|---|---|---|
| DH10B | F– *mcrA* Δ(*mrr-hsdRMS-mcrBC*) φ80*lacZ*ΔM15 Δ*lacX74 recA1 endA1 araD139* Δ (*ara-leu*)7697 *galU galK* λ– *rpsL*($Str^R$) *nupG* | Thermo Scientific |
| Chlamydia trachomatis | Serovar LGV L2 (434) | ATCC® VR-902B™ [8] |
| eHL18 | DH10B expressing pHL18 | This study |
| eHL20 | DH10B expressing pHL20 | This study |
| eHL54 | DH10B expressing pHL54 | This study |
| eHL55 | DH10B expressing pHL55 | This study |
| eHL92 | DH10B expressing pHL92 | This study |
| eHL93 | DH10B expressing pHL93 | This study |
| cHL54 | Chlamydia trachomatis strain expressing Cdu1-mAID-FLAG by its genome | This study |
| cHL55 | Chlamydia trachomatis strain expressing Cdu1-FLAG by its genome | This study |
| cHL92 | Chlamydia trachomatis strain expressing IncA-mAID-FLAG by its genome | This study |
| cHL93 | Chlamydia trachomatis strain expressing IncA-FLAG by its genome | This study |

Table S4. Primers used in this study

| Primer | Sequence 5' to 3' | Description |
|---|---|---|
| 3K_Cdu1_down_F_40 | GATGAACTATACAAGTAAACTCTTTTCTAATCTAAAAATCTTTTTAAATAAGAGG | Inserts Cdu1 3 kb downstream sequence into pKW-L2 plasmid, for pHL54, pHL55 |
| 3K_Cdu1_down_R_40 | TGACGCCCTGCAGGTCTCGTTAATCCTCTTCATAAGGTTTCAG | Inserts Cdu1 3 kb downstream sequence into pKW-L2 plasmid, for pHL54, pHL55 |
| 3K_Cdu1_up_F_40 | GCTTAAAAAGCGGTCGACCAAAATTCTCTCATCTGAAACTATTTGCTAAC | Inserts Cdu1 3 kb upstream sequence into pKW-L2 plasmid, for pHL54, pHL55 |
| 3K_Cdu1_up_R_40 | GCCGGAGCCCCCTCTATTACTCTAGAATCGCAGAGCAATTTCCC | Inserts Cdu1 3 kb upstream sequence into pKW-L2 plasmid, for pHL54, pHL55 |
| 54/55_vector_F_40 | AAACCTTATGAAGAGGATTAACGAGACCTGCAGGGCGTCAGACC | Amplifies the pKW-L2 backbone, for pHL54, pHL55 |
| 54/55_vector_R_40 | AGTTTCAGATGAGAGAATTTTGGTCGACCGCTTTTTAAGCAAAAGAG | Amplifies the pKW-L2 backbone, for pHL54, pHL55 |
| aadA_F_M_40 | ATAAAACGAAAGGCCCAGTCTTCCGACTGAGCCTTTCGTTTTATATCATCATGCCTCCTCTAGAC | Inserts aadA into pKW-L2, for pHL54, pHL55 |
| aadA_R_40 | GATTTTTAGATTAGAAAAGAGTTTACTTGTATAGTTCATCCATGCCATGTG | Inserts aadA into pKW-L2, for pHL54, pHL55 |

| | | |
|---|---|---|
| cdu1_F_40 | AATTGCTCTGCGATTCTAGAGTAATAGAGGGGCTCCGGC | Inserts Cdu1 into pKW-L2, for pHL54, pHL55 |
| cdu1_R_54_M_40 | AACGAAAGGCTCAGTCGGAAGACTGGGCCTTTCGTTTTATTTACTTGTCGTCATCGTCTTTGTAG | For insertion of mAID-fused Cdu1 into pKW-L2, for pHL54 |
| cdu1_R_55_M_40 | AGAGGAGGCATGATGATTTACTTATCGTCGTCATCCTTGTAATC | For insertion of WT Cdu1 into pKW-L2, for pHL55 |
| 92_aada_F | CCTCTACAAAAAGCTCCGGAGGTGGATCGGGAGGTGG | Inserts IncA 3 kb downstream sequence into pKW-L2 plasmid, for pHL92 |
| 92_aada_R | CAAAAAGCTCCTAAGATTACTTGTATAGTTCATCCATGCC | Inserts IncA 3 kb downstream sequence into pKW-L2 plasmid, for pHL92 |
| 92_downstream_F | AACTATACAAGTAATCTTAGGAGCTTTTTGCAATGCAAAAC | Inserts IncA 3 kb upstream sequence into pKW-L2 plasmid, for pHL92 |
| 92_downstream_R | GGTCTGACGCCCTGCAGGTCCTTGATTTGTCTCTGGACCC | Inserts IncA 3 kb upstream sequence into pKW-L2 plasmid, for pHL92 |
| 92_upstream_F | AGCGGTCGACCACAATCAATACTTTCTTCATTAACTAAGCC | Amplifies the pKW-L2 backbone, for pHL92 |
| 92_upstream_R | CCGATCCACCTCCGGAGCTTTTTGTAGAGGGTG | Amplifies the pKW-L2 backbone, for pHL92 |
| 92_vector_F | ACAAATCAAGGACCTGCAGGGCGTCAGACC | Inserts aadA into pKW-L2, for pHL92 |

| | | |
|---|---|---|
| 92_vector_R | TGAAGAAAGTATTGATTGTGGTCGACCGCTTTTTAAGC | Inserts aadA into pKW-L2, for pHL92 |
| 93_F | CCCTCTACAAAAAGCTCCAGCGCTGACTACAAAGACGATG | Removes the mAID from pHL92, to generate pHL93 |
| 93_R | GTAGTCAGCGCTGGAGCTTTTTGTAGAGGGTG | Removes the mAID from pHL92, to generate pHL93 |
| gapA_RT_F | CACACGGATCTTTCGCTCCT | For qRT-PCR that amplifies the glyceraldehyde-3-phosphate dehydrogenase (*gapA*) gene |
| gapA_RT_R | CGACGACGACATCAACATCC | For qRT-PCR that amplifies the *gapA* gene |
| rrs_RT_F | GCGTGGATGAGGCATGCAGT | For qRT-PCR that amplifies the Chlamydial 16S rRNA |
| rrs_RT_R | ACCGTCATCCTGGAGCGTT | For qRT-PCR that amplifies the Chlamydial 16S rRNA |

Table S5. Antibodies used in this study

| Antibodies | Dilution | Description & Source |
|---|---|---|
| anti-cHSP60 | 1:200 (IF)<br>1:50 (ExM) | Primary antibody, Santa Cruz, #sc-57840 |
| anti-p62 | 1:200 (IF) | Primary antibody, Santa Cruz, #sc-28359 |
| anti-OmcB | 1:1000 (WB) | Primary antibody, Invitrogen, #PA5-117552 |
| anti-ompA | 1:500 (WB) | Primary antibody, Invitrogen, #PA5-117609 |
| anti-IncA | 1:100 (WB) | Primary antibody, self-made |
| Anti-DDDDK tag | 1:500 (IF)<br>1:200 (ExM) | Primary antibody, Abcam, #ab205606 |
| Alexa Fluor™ Plus 488 | 1:300 (IF) | Secondary antibody, Invitrogen, #A32723 |
| Alexa Fluor™ Plus 555 | 1:300 (IF) | Secondary antibody, Invitrogen, #A32732 |
| anti-Rabbit IgG (H+L), HRP | 1:50000 (WB) | Secondary antibody, Invitrogen, #31460 |
| anti-Mouse IgG (H+L), HRP | 1:50000 (WB) | Secondary antibody, Invitrogen, #31430 |
| anti-alpha Tubulin HRP | 1:50000 (WB) | Hrp preconjugated antibodies, Abcam, #ab185067 |
| anti-Myc HRP | 1:1000 (WB) | Hrp preconjugated antibodies, Abclonal, #AE026 |
| anti-DDDDK-Tag HRP | 1:5000 (WB) | Hrp preconjugated antibodies, Abclonal, #AE095 |
| anti-GAPDH HRP | 1:25000 (WB) | Hrp preconjugated antibodies, Abclonal, #19056 |

| | | |
|---|---|---|
| anti-β-Actin | 1:25000 (WB) | Hrp preconjugated antibodies, Abclonal, # AC028 |

[a]ExM (Expansion Microscopy), IF (Immunofluorescence), WB (Western Blot)

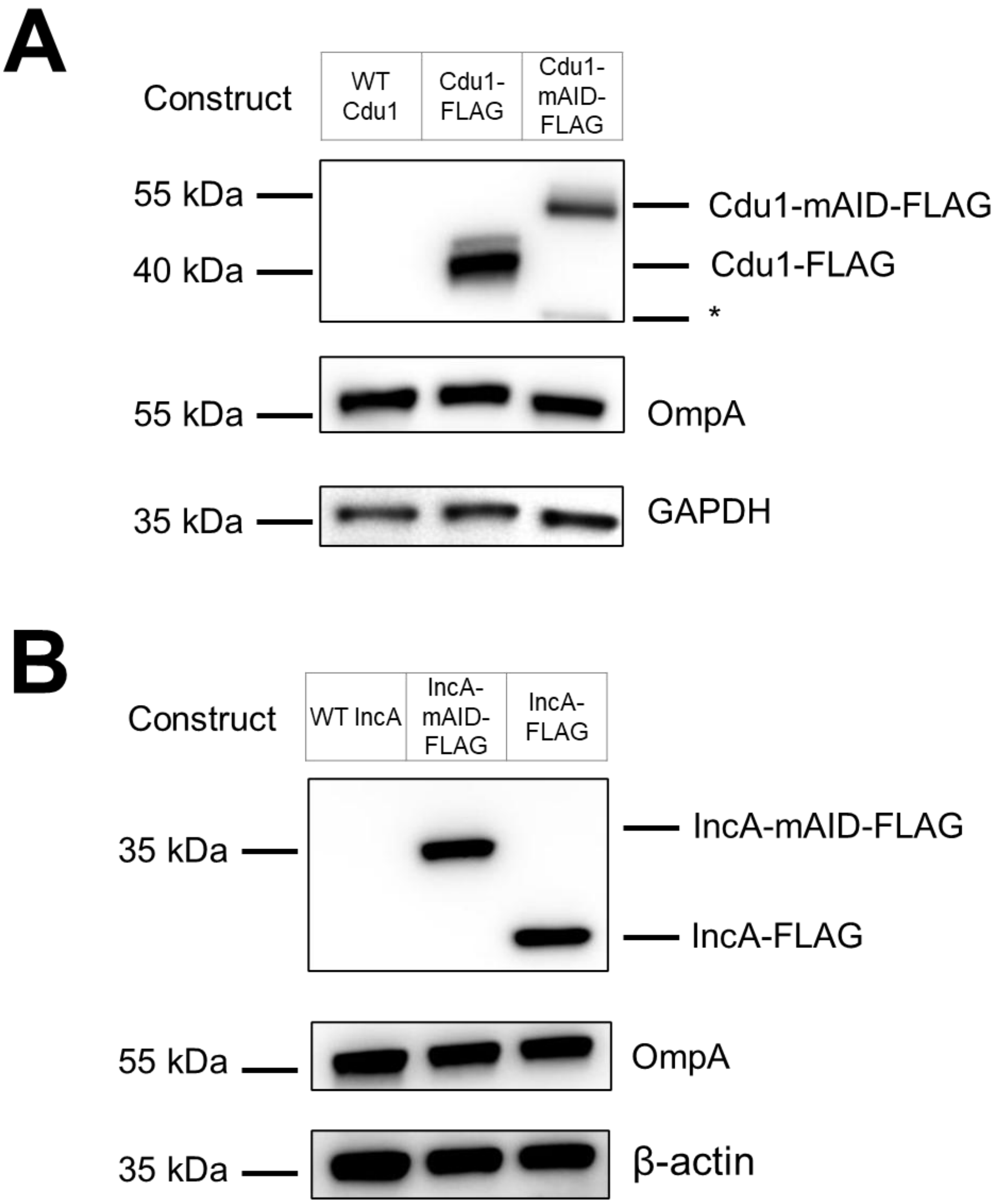


**Figure S1**. (A) Immunoblot analysis of Cdu1-FLAG and Cdu1-mAID-FLAG expression in recombinant chlamydial strains. Cdu1 constructs were detected with anti-FLAG antibody; *Chlamydia* OmpA and host cell GAPDH served as a loading control. (B) Immunoblot analysis of IncA-FLAG and IncA-mAID-FLAG expression in recombinant strains. IncA constructs were detected with anti-FLAG antibody; *Chlamydia* OmpA and host β-actin served as loading control.

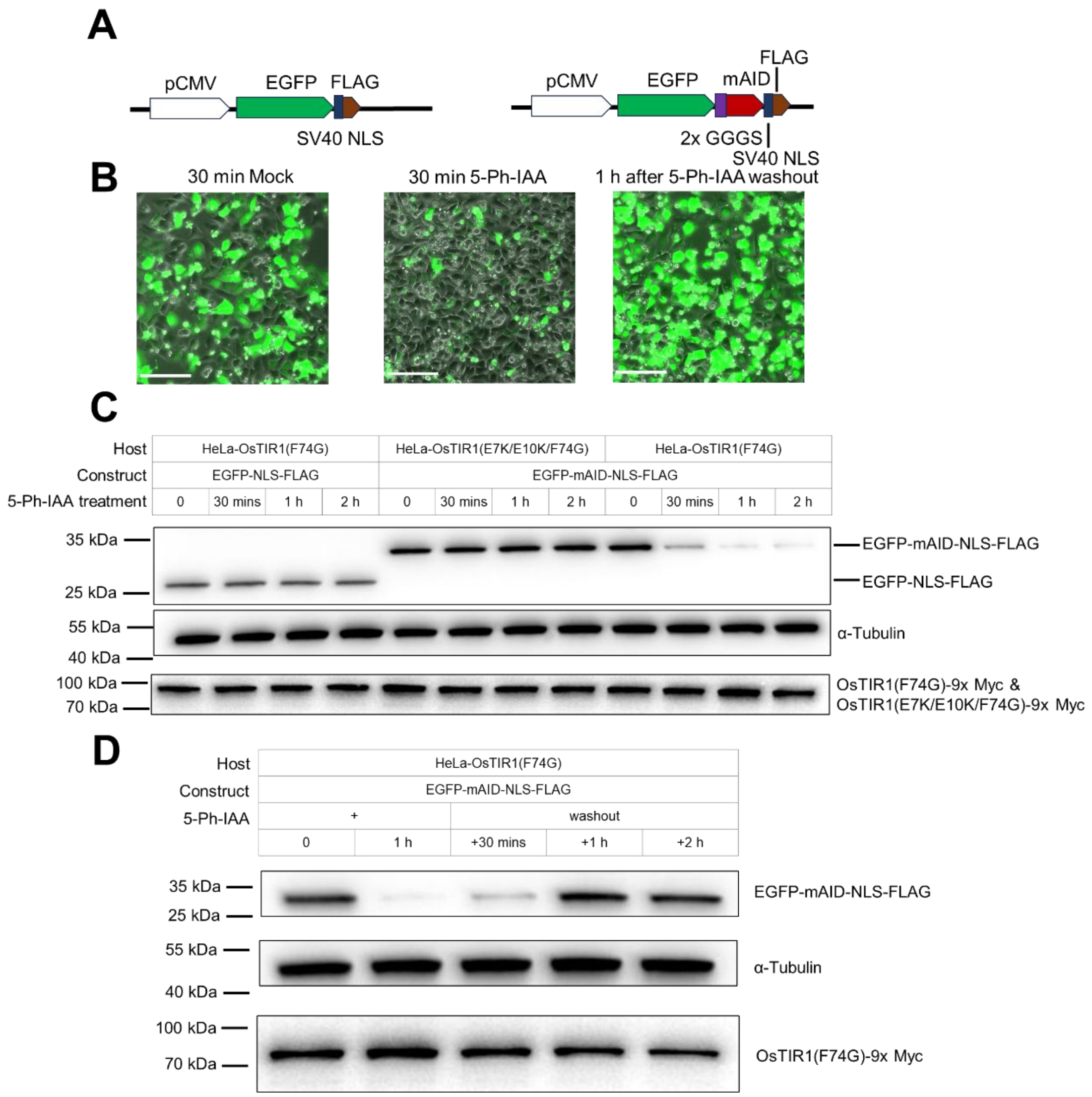


**Figure S2. Development and validation of the AID2 system for spatiotemporal control of protein abundance in HeLa cells.** (A) Schematic of cassette expressing the GFP mutants used for degradation. SV40 NLS, nuclear localization signal. (B) Time-resolved GFP depletion and recovery. Live-cell microscopy of HeLa cells co-expressing OsTIR1(F74G) and mAID-GFP. GFP fluorescence diminishes within 30 minutes of 1 μM 5-Ph-IAA treatment and recovers 1 hour after auxin washout. Scale bar = 100 μm. (C) Immunoblot validation of AID2 specificity. Robust GFP degradation requires both mAID tagging and functional OsTIR1(F74G). Minimal degradation occurs with untagged GFP or catalytically impaired OsTIR1(E7K/E10K/F74G). α-tubulin serves as a loading control; OsTIR1-F74G expression was detected using anti-Myc antibody. (D) Reversibility of AID2-mediated degradation. Cells were treated with 1 μM 5-Ph-IAA for 30 minutes, after which auxin was removed and GFP levels were assessed at the indicated recovery time points. GFP levels rebound within 1 hour of auxin removal, demonstrating dynamic control. All experiments were replicated ≥3 times with consistent results.

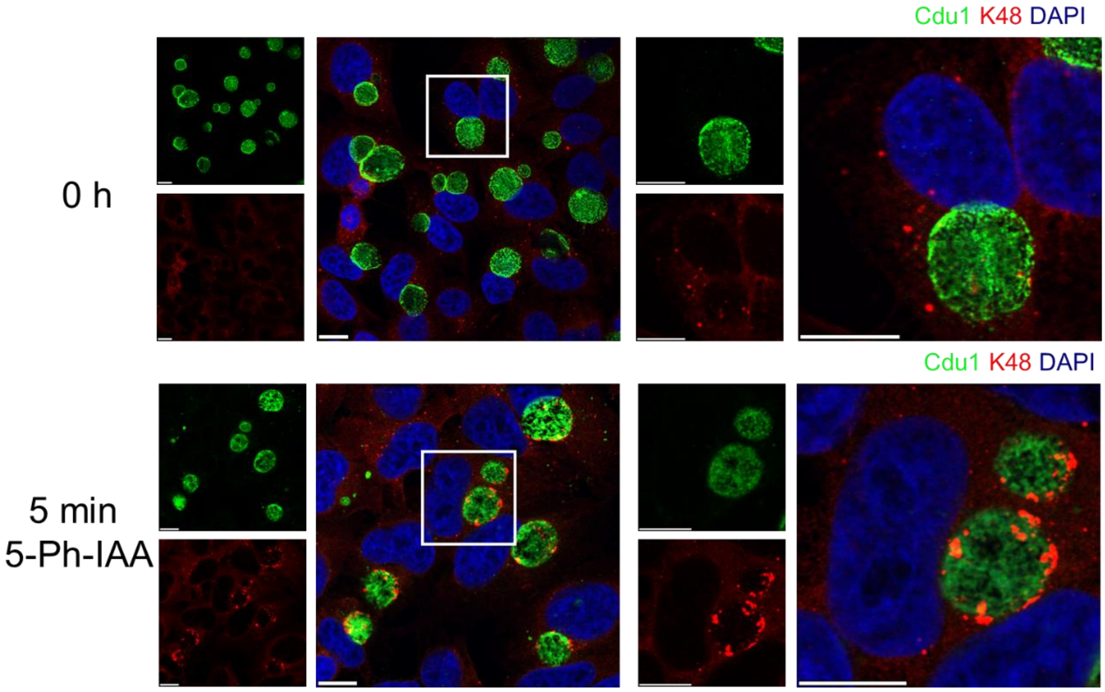


**Figure S3. Ctr-AIDE induces rapid K48-ubiquitin conjugation to Cdu1 during degradation.** Immunofluorescence microscopy reveals K48-ubiquitin recruitment to the inclusion membrane within 5 minutes of auxin treatment. The rapid K48-ubiquitin signal reflecting host-mediated tagging of Cdu1-mAID during degradation, may interfere with the downstream signal caused by inclusion ubiquitination due to Cdu1 degradation. Cdu1 (green, anti-FLAG) was visualized with K48-Ubiquitin (red), and DAPI (blue) marking bacterial and host DNA. Scale bar = 10 µm.

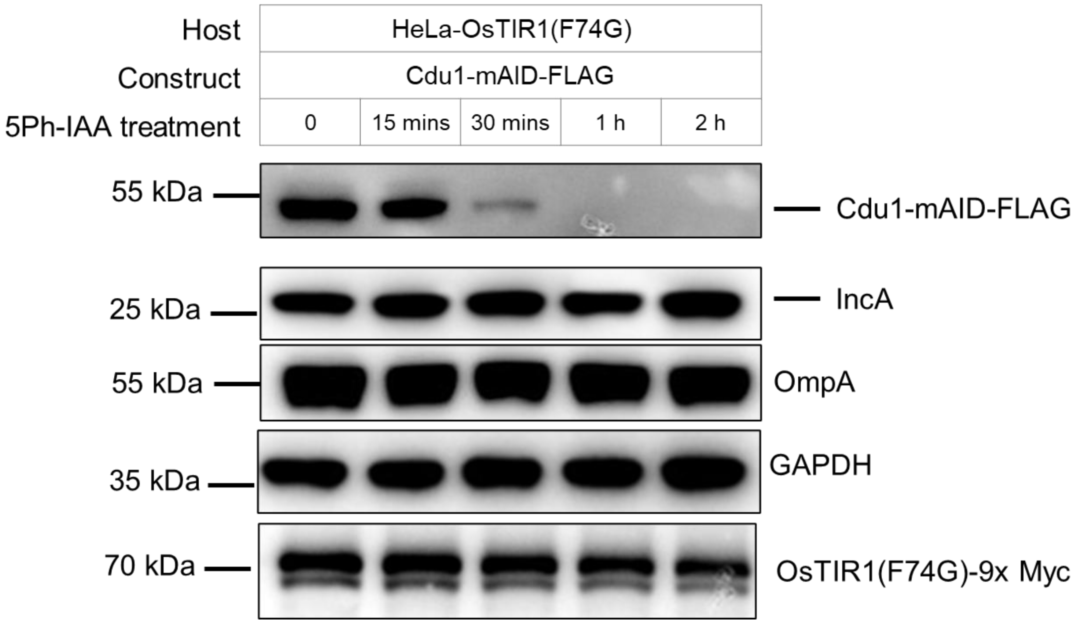


**Figure S4. Ctr-AIDE selectively degrades mAID tagged effector.** HeLa cells expressing OsTIR1(F47G) were infected with Cdu1-mAID expressing strains at MOI=1. Cdu1 (anti-FLAG), IncA (anti-IncA) and OsTIR1 mutants (anti-Myc) levels were monitored, with Chlamydial Major Outer Membrane Protein (OmpA) as a loading control and GAPDH as a host cell control.

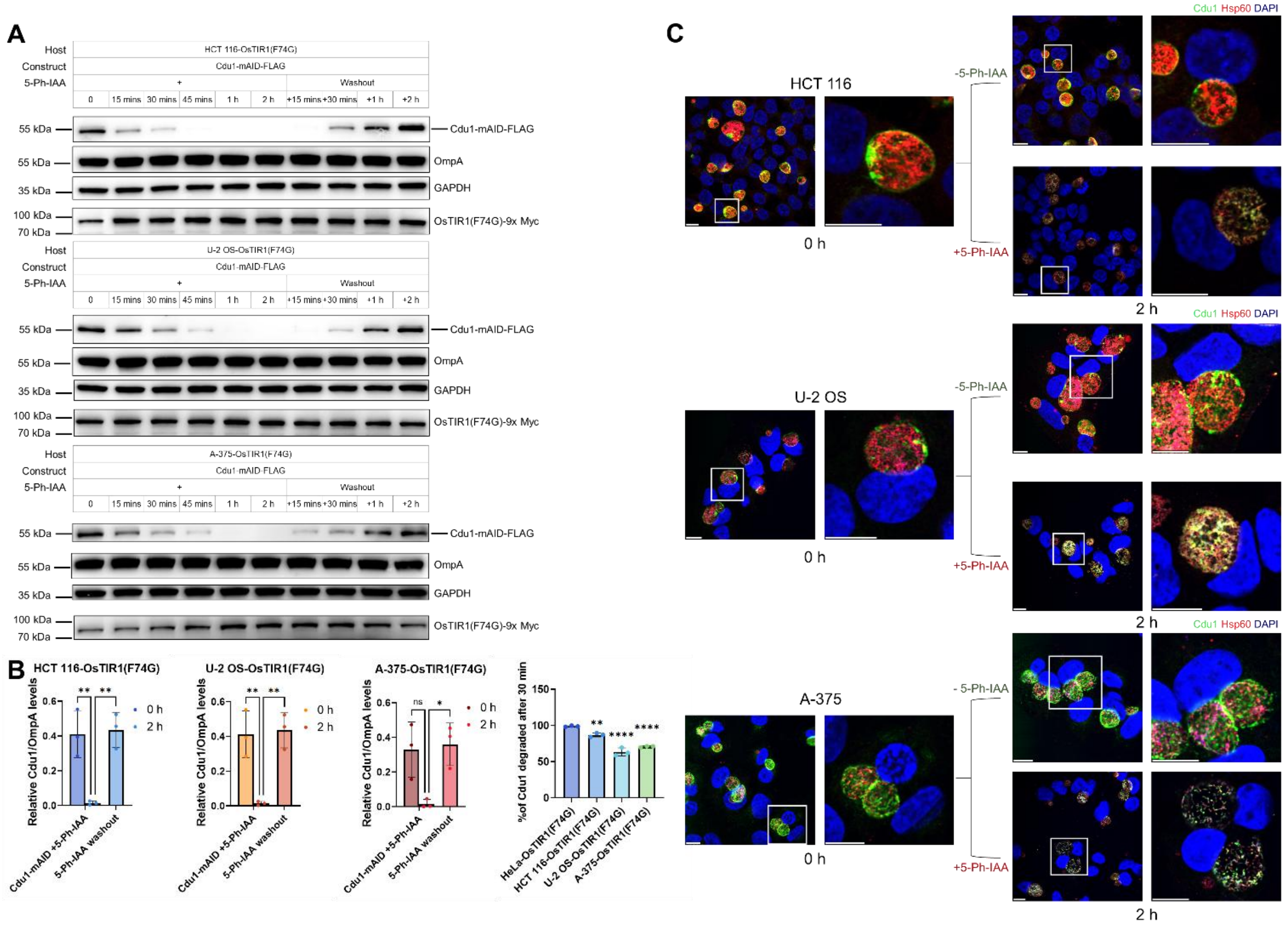


**Figure S5. Ctr-AIDE-mediated Cdu1 degradation in various cancer cell lines.** (A) Ctr-AIDE-mediated regulation of Cdu1-mAID expression in different cancer cell lines. A-375, HCT 116 and U-2 OS cells expressing OsTIR1(F74G) were infected with Cdu1-mAID expressing *Chlamydia* (MOI = 1). Cells were treated with 1 µM 5-Ph-IAA for 2 hours (degradation) or further washed and incubated in 5-Ph-IAA free medium for 2 hours (recovery). Cdu1 (anti-FLAG) and OsTIR1(F74G) (anti-Myc) levels were monitored, with *Chlamydia* OmpA as a loading control and host GAPDH as a cellular control. (B) Quantification and comparation of Cdu1 degradation efficiency in various cancer cell lines. Quantification (left three panels) shows Cdu1-FLAG levels normalized to OmpA (mean ± SD; n = 3 biological replicates). Significance assessed by two-tailed paired t-test (**p < 0.01; *p < 0.05; n.s., not significant). Comparation (right panel) shows relative decrease of Cdu1 levels after 30 minutes of 1 µM 5-Ph-IAA treatment (normalized to 0-hour controls). Data represent mean percentage decrease ± SD (n = 3 biological replicates; one representative blot in Panel A. Significance versus HeLa: ****p < 0.0001, **p < 0.01 (one-way ANOVA with Tukey Multiple comparisons test). (C) Immunofluorescence microscopy confirms Cdu1 depletion dynamics in A-375, HCT 116 and U-2 OS cells. Cdu1 (green, anti-FLAG) was visualized with *Chlamydia* Hsp60 (red), to mark inclusions and DAPI (blue) to label bacterial and host DNA. Scale bar = 10 µm. All experiments were replicated ≥3 times with consistent results.

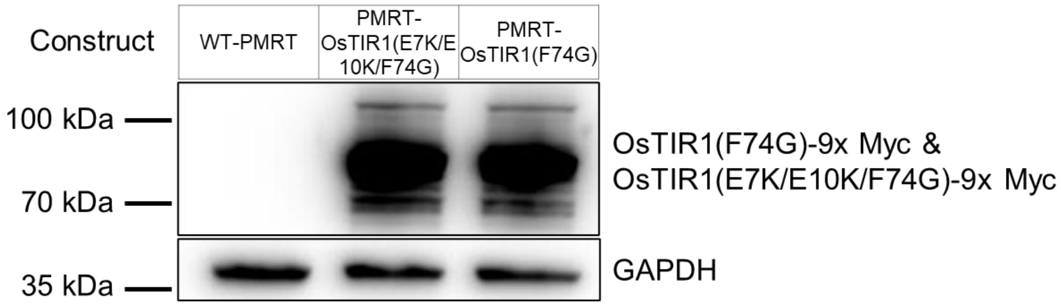


**Figure S6. Expression of OsTIR1(F74G) and OsTIR1(E7K/E10K/F74G) in PMRT cells.** OsTIR1 mutants (anti-Myc) were monitored, with GAPDH as a loading control.

.

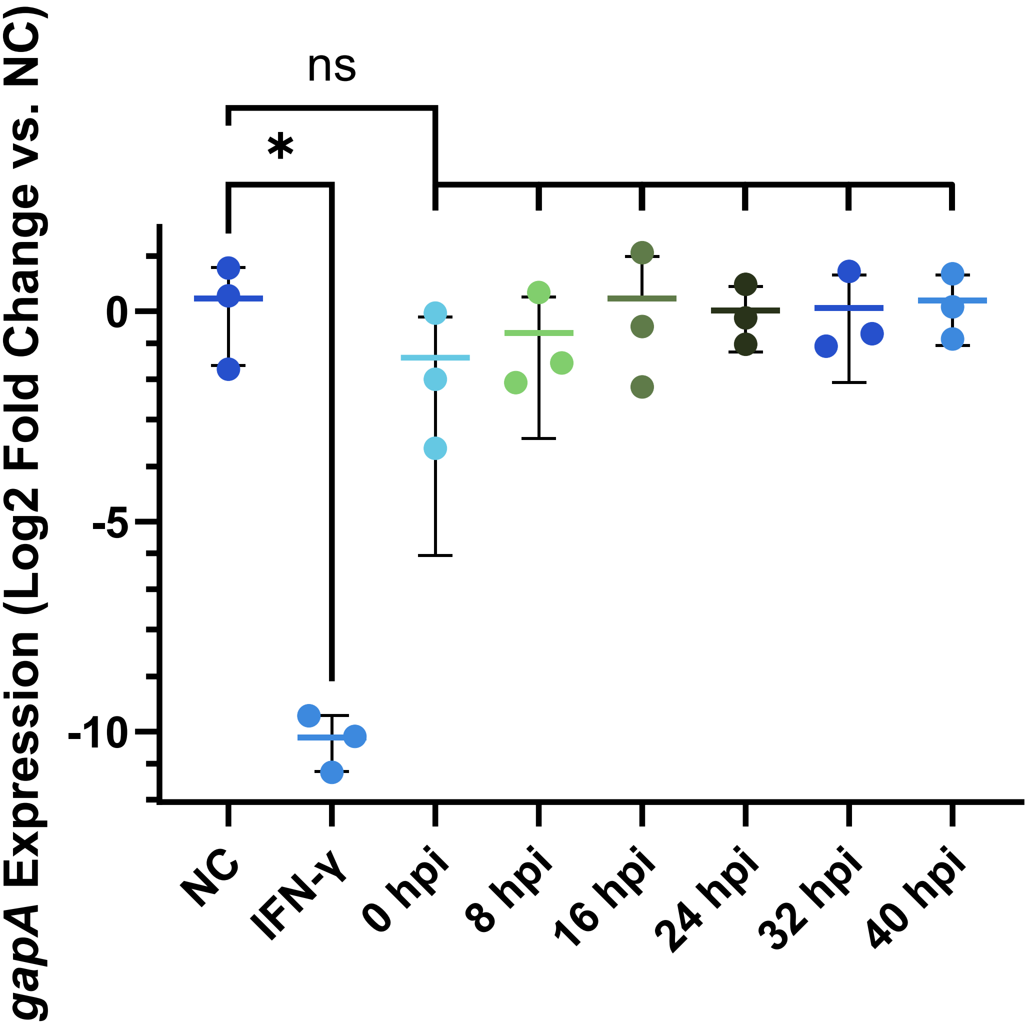


**Figure S7. Cdu1 depletion does not impair *Chlamydia* metabolic activity in HeLa cells**. RT-qPCR analysis of *gapA* (glyceraldehyde-3-phosphate dehydrogenase, metabolic marker) and *rrs* (16S rRNA, load control) in Cdu1-mAID strains treated with 5-Ph-IAA at indicated timepoints to 48 hpi. IFN-γ (40-hour treatment, 50 U/mL) served as a positive control for metabolic inhibition. Data normalized to untreated controls. Significance assessed by one-way ANOVA with Dunnett Multiple comparisons test (*p < 0.05; n.s., not significant).

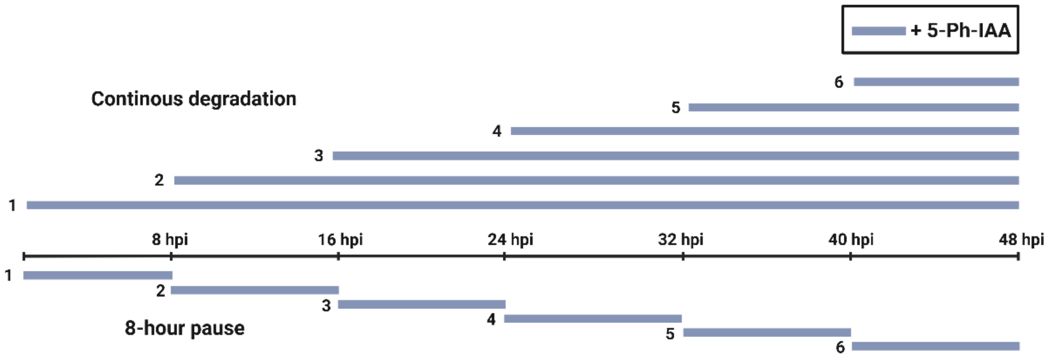


**Figure S8**. **Schematic of temporal Ctr-AIDE-mediated Cdu1 degradation to evaluate its role in chlamydial growth.** The blue lines indicate periods of 1 µM 5-Ph-IAA treatment to induce Cdu1-mAID degradation. Bacterial growth and progeny infectivity were assessed at 48 hpi.

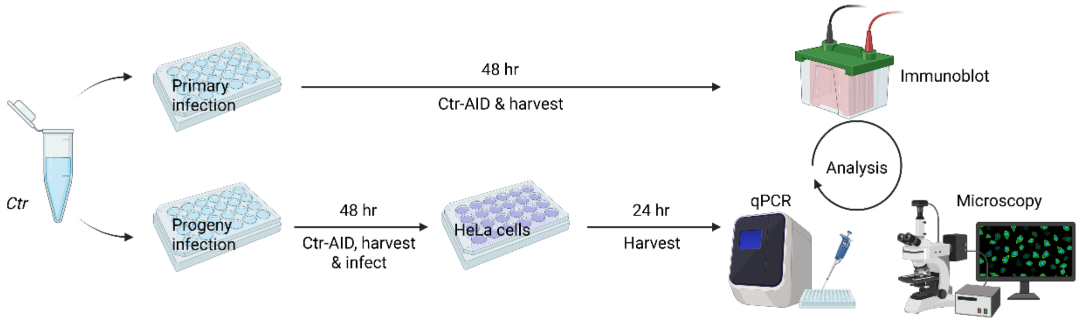


**Figure S9. Schematic representation of the primary and progeny infection assays.**

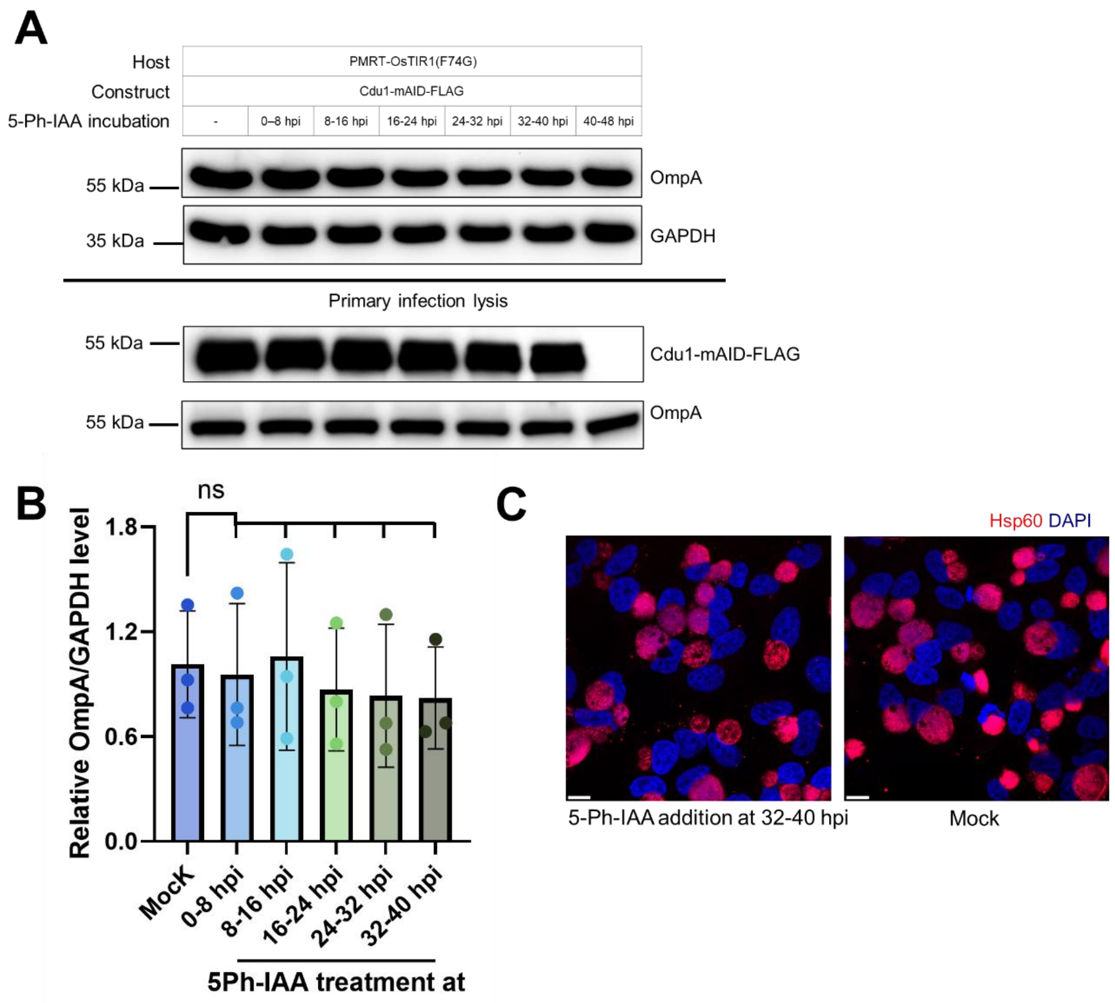


**Figure S10. Short-term Cdu1 depletion does not affect *C. trachomatis* progeny infectivity in primary cells.** (A) Chlamydial lysates from primary infection (MOI=1, Cdu1 degraded at the indicated time points for 8 hours) were normalized by OmpA levels ('Primary infection lysis' part of the immunoblot image) and used to infect new batch of HeLa cells. 24 hpi, OmpA (Progeny activity) and GAPDH (loading control) were analyzed via immunoblot. (B) Quantification of effects of 8 hours Cdu1 expression pause on progeny infectivity. OmpA levels were calculated (normalized to GAPDH, mean ± SD) from three replicates (one representative in Panel A). Statistical significance determined by one-way ANOVA with Dunnett Multiple comparisons test (n.s., not significant). (C) Immunofluorescence microscopy reveals unimpaired progeny formation. HeLa cells were infected with progeny derived from Cdu1-degraded (degraded from 32 to 40 hpi) or control *Chlamydia*. Images were captured at 24 hpi. Inclusions marked by Hsp60 (red) and DNA (DAPI, blue) marking bacterial and host nuclei; scale bar = 10 µm. All experiments were replicated ≥3 times with consistent results.

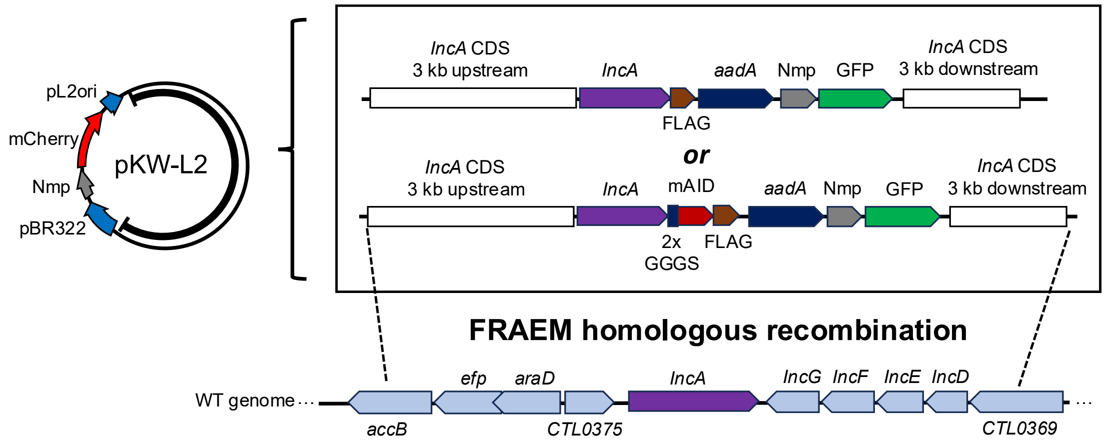


**Figure S11**. Plasmids used in FRAEM homologous recombination for IncA.

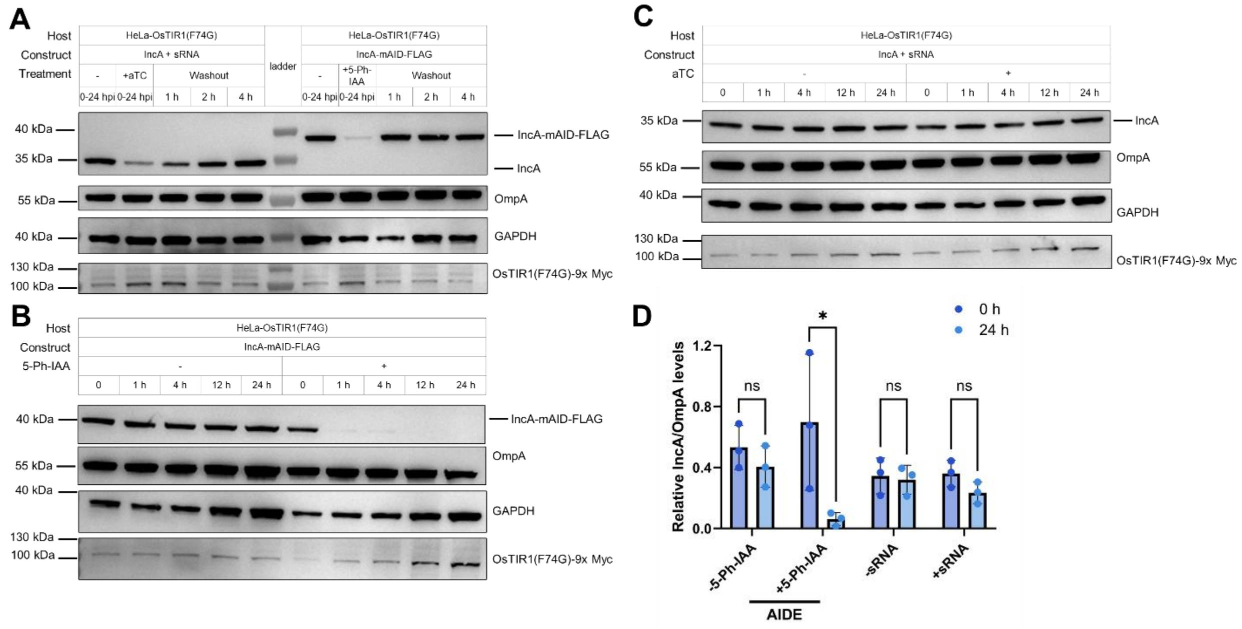


**Figure S12. Comparison of Ctr-AIDE-mediated and sRNA-based IncA depletion in *C. trachomatis*.** (A) Comparison of sRNA silencing and Ctr-AIDE-mediated regulation of IncA expression from 0–24 hpi. Both approaches reduced IncA levels. HeLa cells expressing OsTIR1(F74G) were infected with *C. trachomatis* expressing Cdu1-mAID (MOI = 1). Cells were treated with 1 µM 5-Ph-IAA or 200 ng/mL anhydrotetracycline (aTC; to induce sRNA expression), then washed and incubated in inducer-free medium for the indicated times. Cdu1 (anti-FLAG) and OsTIR1(F74G) (anti-Myc) levels were monitored, with *Chlamydia* OmpA as a loading control and host GAPDH as a cellular control. (B) Ctr-AIDE enabled rapid and efficient depletion of IncA. At 24 hpi, cells were treated with 1 µM 5-Ph-IAA or DMSO for the indicated times. (C) sRNA silencing initiated at 24 hpi failed to deplete IncA. At 24 hpi, cells were treated with 200 ng/mL aTC (to induce sRNA expression) or DMSO for the indicated times. All experiments were replicated ≥3 times with consistent results. (D) Quantification and comparation of IncA degradation efficiency by Ctr-AIDE and sRNA silencing. Quantification shows IncA levels normalized to OmpA (mean ± SD; n = 3 biological replicates). Significance assessed by two-tailed paired t-test (*$p < 0.05$; n.s., not significant).
